# The optimal spatial arrangement of ON and OFF receptive fields

**DOI:** 10.1101/2021.03.10.434612

**Authors:** Na Young Jun, Greg Field, John Pearson

**Affiliations:** Department of Neurobiology, Duke University; Department of Biostatistics & Bioinformatics, Duke University

## Abstract

Many sensory systems utilize parallel ON and OFF pathways that signal stimulus increments and decrements, respectively. These pathways consist of ensembles or grids of ON and OFF detectors spanning sensory space. Yet encoding by opponent pathways raises a question: How should grids of ON and OFF detectors be arranged to optimally encode natural stimuli? We investigated this question using a model of the retina guided by efficient coding theory. Specifically, we optimized spatial receptive fields and contrast response functions to encode natural images given noise and constrained firing rates. We find that the optimal arrangement of ON and OFF receptive fields exhibits a transition between aligned and anti-aligned grids. The preferred phase depends on detector noise and the statistical structure of the natural stimuli. These results reveal that noise and stimulus statistics produce qualitative shifts in neural coding strategies and provide novel theoretical predictions for the configuration of opponent pathways in the nervous system.

**Significance Statement:** Across a wide variety of species, cells in the retina specialized for signaling either increases (ON) or decreases (OFF) in light represent one of the most basic building blocks of visual computation. These cells coordinate to form mosaics, with each cell responsible for a small, minimally-overlapping portion of visual space, but the ways in which these mosaics could be spatially coordinated with each other are relatively unknown. Here, we show how efficient coding theory, which hypothesizes that the nervous system minimizes the amount of redundant information it encodes, can predict the relative spatial arrangement of ON and OFF mosaics. The most information-efficient arrangements are determined both by levels of noise in the system and the statistics of natural images.

## INTRODUCTION

Across many sensory systems, neurons encode information about either increments or decrements of stimuli in the environment: so-called ‘ON’ and ‘OFF’ signals. This division between ON and OFF signaling has been observed in visual Joesch *et al.* (2010), Kuffler (1953), thermosensory Gallio *et al.* (2011), auditory Smith and Lewicki (2006), olfactory Chalasani *et al.* (2007), and electro-sensory Bell (1990) systems. This organization has the advantage that neurons can be tasked with signaling increments or decrements in steady-state stimulus levels with fewer spikes, thereby resulting in more efficient neural codes Barlow (1961), Gjorgjieva *et al.* (2014). Moreover, when the number of potential stimuli is large, neurons often specialize: for example, they only respond to a small region of visual space or a narrow auditory frequency band. The combination of these coding strategies raises two questions: First, how should a particular set of detectors, either the ON and OFF cells, arrange themselves most efficiently to cover stimulus space? Second, what is the optimal relative arrangement of ON and OFF detector grids? For one system, the retina, the answer to the first question is clear from previous work: detectors of a particular type tile stimulus space and exhibit overlap near the 1-sigma boundary of a Gaussian receptive field Devries and Baylor (1997), Borghuis *et al.* (2008), Doi *et al.* (2012), Gauthier *et al.* (2009a,b). The answer to the second question, what might be called the “sensor alignment problem,” has received comparatively little attention and is the focus of this study.

Conceptually, there are three general possibilities for how the sensor alignment problem could be solved: One possibility is that the grids of sensors are statistically independent, meaning the locations of receptive fields in one grid provide no information about the receptive field locations in the other grid. A second possibility is that the two grids are aligned, meaning the receptive field centers in one grid are closer than expected by chance. The third possibility is that the two grids are anti-aligned, meaning the receptive field centers in the two grids are further apart than expected by chance. On general information theory grounds, the optimal solution is likely to depend on noise in the encoding process and the statistics of the encoded stimuli Atick and Redlich (1990, 1992).

While most anatomical studies of retinal mosaics indicate they are statistically independent (16–18, but see 19), we have recently shown that grids of ON and OFF receptive field (henceforth called ‘mosaics’) formed by retinal ganglion cells (RGCs) are anti-aligned when those cells encode similar visual features Roy *et al.* (2021). Here, we show how these results can be explained through the lens of efficient coding theory Barlow (1961). This theory argues that sensory systems should aim to reduce the redundancy present in sensory input while minimizing metabolic costs, thereby reliably encoding natural stimuli with fewer spikes. Efficient coding theory has been successful at explaining many aspects of sensory processing and retinal physiology, including center-surround receptive fields, the formation of mosaics, and a greater proportion of OFF than ON cells Barlow (1961), Doi *et al.* (2012), Atick and Redlich (1992), Karklin and Simoncelli (2011), Ratliff *et al.* (2009). Thus, we asked whether efficient coding theory might predict the optimal spatial arrangement of ON and OFF receptive field mosaics within the retina. Our approach to this question involved optimizing a model that approximates the processing performed by many RGCs Karklin and Simoncelli (2011). By maximizing the mutual information between an (input) library of natural images and (output) spike rates, we examined the effects of image statistics and encoding noise on the optimal arrangement of ON and OFF mosaics.

In this model, we found that the optimal spatial arrangement was a pair of approximately hexagonal mosaics of ON and OFF receptive fields. However, surprisingly, the relative alignment of these mosaics depended on the input noise, output noise, and the statistics of the natural image set. When output noise was low, the mosaics were aligned, with ON and OFF receptive fields centered at nearby locations more often than expected by chance. When output noise was relatively high, anti-alignment became the favored arrangement. Surprisingly, the content of the image set also strongly influenced the transition between aligned and anti-aligned mosaics. In particular, when image sets contained more ‘outlier’ images with particularly large luminance or contrast values, anti-alignment became the favored state for fixed input and output noise. We demonstrate analytically and confirm computationally that as noise parameters or stimulus statistics vary, mutual information changes smoothly, while the optimal mosaic arrangements undergo a sudden, qualitative shift. Finally, we confirm these predictions by showing that systematic manipulations of the training data set change the phase boundary in a manner predicted by an analytical model. These findings underscore the crucial role played by both noise and the statistics of natural stimuli for understanding specialization and coordination in sensory processing.

## METHODS

### Linear-nonlinear efficient coding model

The model used to examine efficient coding is based on previous work Karklin and Simoncelli (2011), designed to find the optimal encoding of visual scenes. The model takes patches of natural scenes Doi and Lewicki (2007) as the input, each of which has 18 × 18 (324) pixels. Each input image was multiplied by a circular mask to avoid cases in which corners in the optimization region produce corner artifacts that dominate the final configuration. The circular masking effectively left about 255 pixels in each input image patch. The model consisted of linear-nonlinear (LN) architecture with learned linear filters and nonlinearities. Given an input stimulus **x**, an input noise **n**_*x*_, and the linear filter **w**_*j*_ for each neuron *j*, the nonlinearities were modeled as softplus functions of the linearly filtered noisy stimulus for each neuron 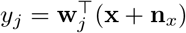:

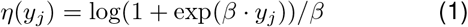

using a fixed value of *β* = 2.5. We empirically chose a fixed value of *β* = 2.5 which produced a stable optimization trajectory. The output of the model consisted of 100 units that mimic retinal ganglion cells (RGCs), whose firing rates are modeled using neuron-specific gain and threshold parameters *γ_j_* and *θ_j_*, plus an additive output noise *n_r,j_*:

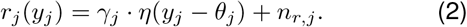

Note that scaling *r, γ*, and *n_r_* by the same factor leaves the model invariant, so we choose units in which the mean firing rate is 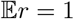. We used a first-order approximation that treats the conditional distribution of activation as a Gaussian, which allowed us to derive a closed-form expression for the mutual information. A detailed derivation is provided in the supplement. The model was trained to maximize the mutual information between the input images and the firing rates of the output neurons. To model the metabolic cost of spiking, the average firing rate of each neuron across images was constrained to 1, which was enforced via an augmented Lagrangian method with quadratic penalty. Input and output noise were modeled as i.i.d. Gaussian noise (zero means, and standard deviations of *σ*_in_ and *σ*_out_ respectively).

The parameters of the model were the receptive field filter weights,the gain of the nonlinearity, and the nonlinearity threshold of each output RGC. To preserve positive gain for each output unit, we optimized the logarithms of the corresponding parameters. Filter weights were initialized by drawing samples from a Gaussian distribution (mean zero, standard deviation 1; see **Fig 1B**, top row). The parameters of the nonlinearities (log gain and threshold) were initialized by sampling from a uniform distribution [0, 1]. The parameters were then optimized using stochastic gradient descent with a learning rate of 0.001. During training, we calculated gradients using mini-batches of 100 image patches at a time, renormalizing the filters to have unit norm after each gradient step. Models were trained for 1,000,000 iterations, which we found sufficient in practice to ensure convergence of the mutual information. As a means of escaping local optima, between 200,000 and 500,000 iterations, two modifications were applied to the optimization procedure, which we call jittering and centering. These were used to speed convergence: The jittering operation was applied every 5,000 iterations and consisted of raising each element of the kernels to the power 1.25; this makes high-amplitude portions of the filters more pronounced while attenuating low-amplitude portions. The centering operation penalizes the spatial spread of the kernel by adding the mean of the spatial variance of each kernel as an additional loss term; this encourages the kernels to be localized around their centers. These two operations allowed kernels that were trying to ‘squeeze’ into an already formed mosaic the space to do so, effectively speeding the optimization and allowing all kernels to contribute to the encoding. After these operations, the model was optimized without the modifications for another 500,000 iterations. We confirmed that jittering and centering do not alter the final converged shape of the kernels.

**FIG. 1.**
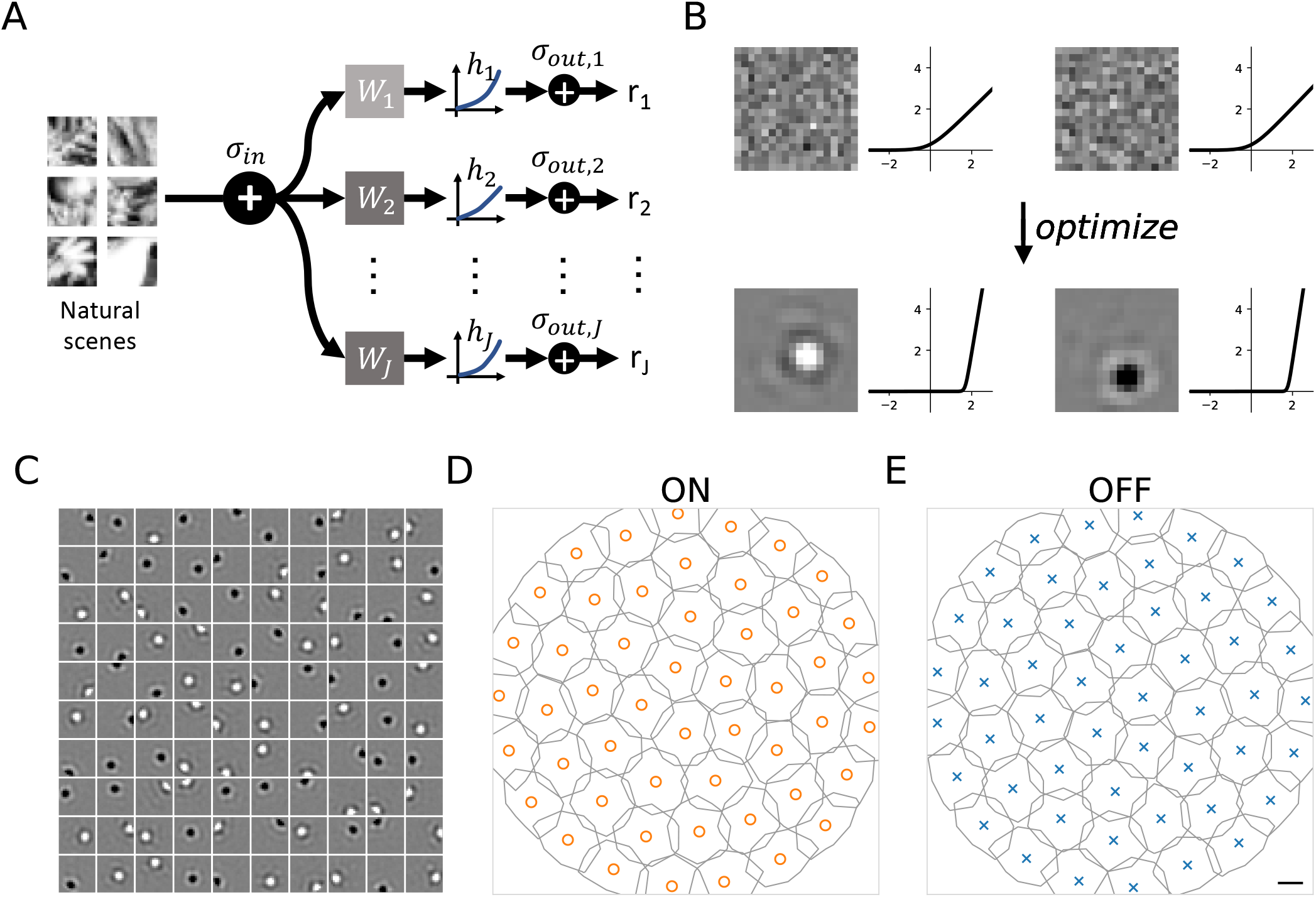
Spatial tiling of receptive fields is predicted by efficient coding. **A.** Efficient coding model architecture. Images of natural scenes (plus input noise *σ*_in_) are multiplied by linear filters **w**_*j*_, passed through a nonlinear function *h_j_*, and perturbed by output noise *σ*_out_, resulting in a firing rate *r_j_* for neuron *j*. **B.** Examples of initial and optimized filters and nonlinearities, where the filters are initialized from a white noise distribution and converge to ON and OFF kernels with a center-surround pattern. The nonlinearities are initialized as unscaled (but slightly perturbed) softplus functions and converge to nonzero threshold values. **C.** A plot of 100 kernels from a trained model with *σ*_in_ = 0.4, *σ*_out_ = 3.0. **D.** A contour plot showing the tiling of the ON and OFF kernels. The contours of the two types of kernels are drawn where the normalized pixel intensity is ± 0.21. Orange circles and blue x’s indicate receptive field centers of mass for ON and OFF cells, respectively. Scale bar is width of one image pixel.

### Median nearest neighbor analysis of the spatial relationships between the mosaics

To quantify the relative alignment of ON and OFF mosaics, we used a metric based on the median nearest-neighbor distances between heterotypic receptive field centers. For each receptive field center, we found the nearest center with the opposite polarity to define its heterotypic nearest neighbor distance, and calculated the median of these values(See Equation 3). When ON and OFF mosaics are nearly aligned, this value is close to zero as in **Fig 2A-B**.

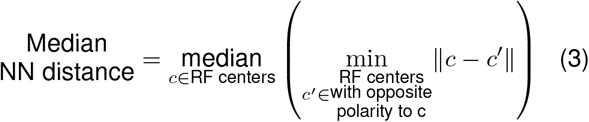

**FIG. 2.**
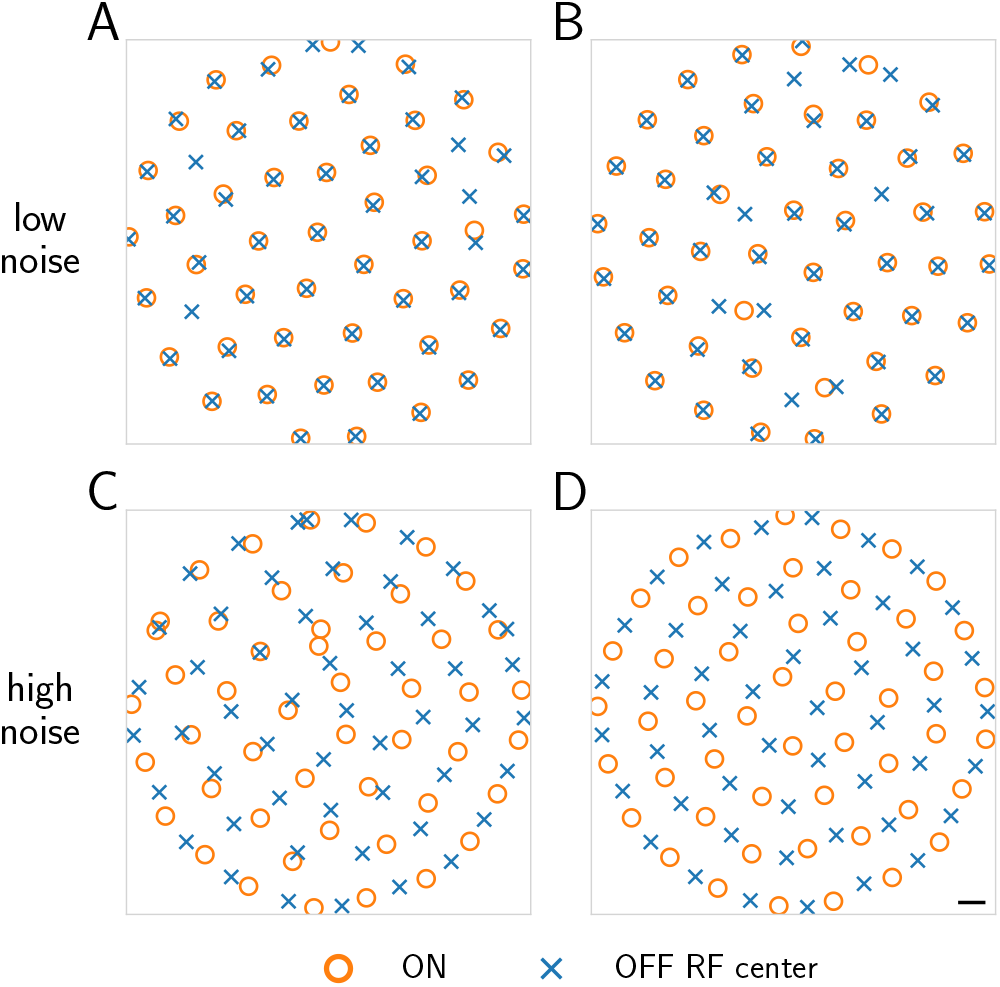
Spatial coordination of receptive field mosaics depends on input and output noise level. Receptive field centers for ON (orange circle) and OFF (blue x) cells under differing sets of noise parameters. **A:** (*σ*_in_=0.02, *σ*_out_=1.0), **B:** (*σ*_in_=0.1, *σ*_out_=1.0), **C:** (*σ*_in_=0.4, *σ*_out_ =2.0), **D:** (*σ*_in_=0.4, *σ*_out_=3.0). The first two parameter sets result in aligned mosaics, while the latter two, at higher levels of output noise, are anti-aligned. Scale bar is width of one image pixel.

### One-shape model for rapid exploration of mosaic arrangements

The full 18×18 system required many hours to optimize, motivating the development of a smaller, scaled-down system for exploring how mosaic arrangements depended on different noise parameters (**Fig 3)** or different stimulus statistics (**Fig 6)**. Here, we restricted input images to 7×7, fixed the number of ON and OFF cells at 7 each (14 total), and adopted a fixed parametric form for the receptive field. Thus, the optimization only learned receptive field center locations and a small number of receptive field shape parameters. The receptive field function was parameterized as a difference of two Gaussian curves, with a third parameter determining their relative magnitude.

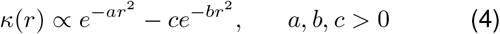

**FIG. 3.**
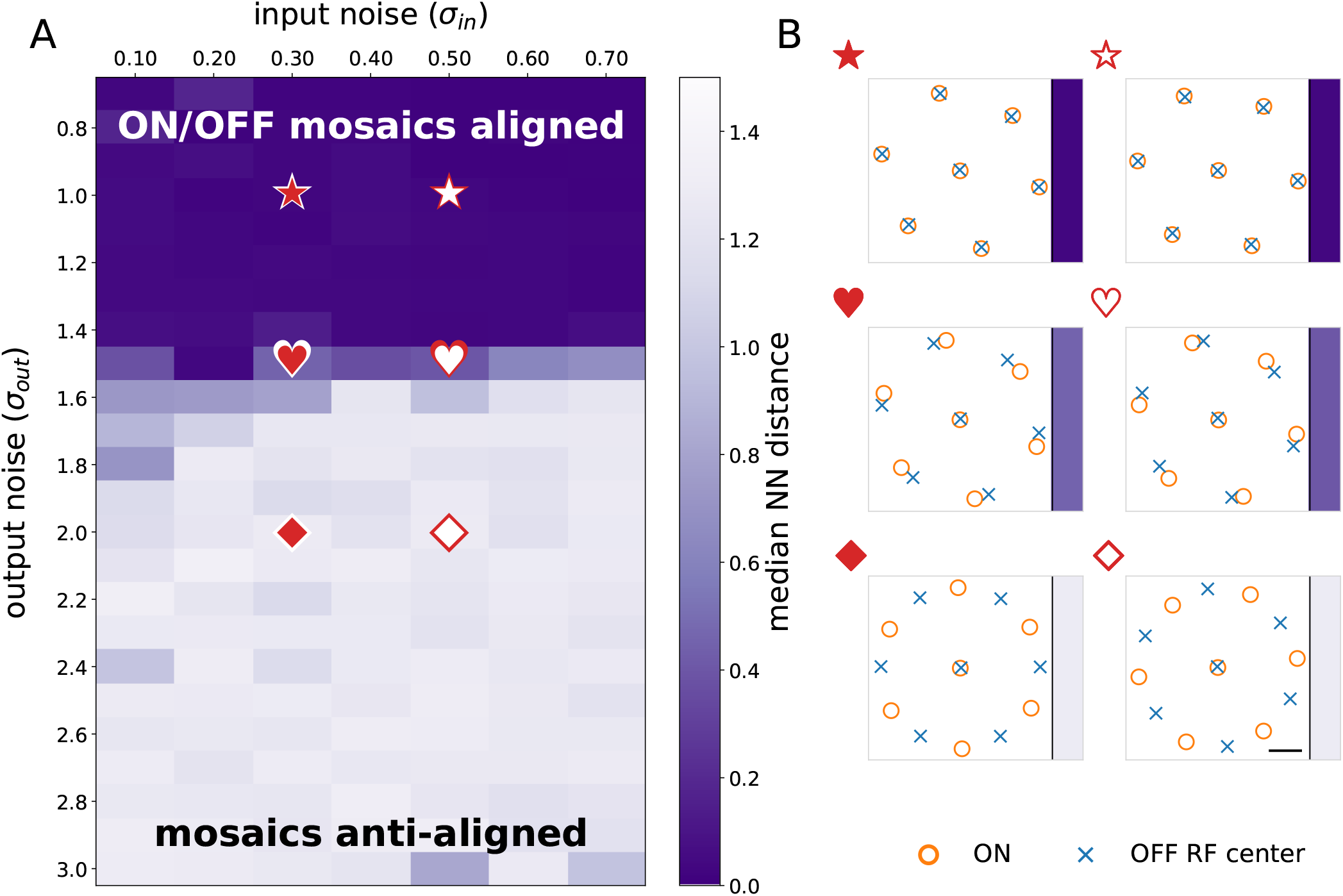
Optimal mosaic configurations transition from aligned to anti-aligned as a function of noise. **A.** As output noise increases, optimal mosaic configurations shift from aligned to anti-aligned, and this holds over a range of input noise values. Color indicates median NN distance, with darker indicating more clearly aligned (see Equation 3 and Methods). The existence of a phase boundary between the two arrangements is clear. **B.** Examples of optimal mosaic arrangements for representative input and output noise combinations. Symbols denote the corresponding locations in 3A. Note the existence of optimal configurations between aligned and fully anti-aligned (hearts) for some parameter values. Scale bar is width of one image pixel.

Optimization in this setup involves fewer parameters, as well as allowing us to use more efficient first-order optimization methods like Adam Kingma and Ba (2014). To examine the dependence of mosaic arrangement on noise a grid search was performed over combinations of input and output noise values, with input noise (*σ*_in_) ranging from 0.0 to 0.75 and output noise (*σ*_out_) from 0.75 to 3.0. As previously argued Karklin and Simoncelli (2011), these values lie within the physiological range for naturalistic stimuli. The optimization was run three times for every pair of noise values and the model resulting in the highest mutual information was retained **Fig 3**. This model was also used to examine the dependence of mosaic alignment on the statistics of natural scenes **Fig 6**. In these simulations the input noise was held fixed while the output noise and distribution of natural images was varied to examine the interaction between these factors on the resulting mosaic arrangements.

## RESULTS

### Linear-nonlinear efficient coding model of the retina

We asked how mosaics of ON and OFF cells should be arranged to efficiently encode natural scenes. To answer this question, we trained a model of retinal processing to maximize the mutual information between a library of natural images and a set of output firing rates (**Fig 1A)**. The model consists of a single layer of linear-nonlinear units intended to approximate the encoding performed by RGCs Borghuis *et al.* (2008), Doi *et al.* (2012), Karklin and Simoncelli (2011), Meister and Berry (1999), Chichilnisky (2001), Baccus and Meister (2002). These model cells linearly filter the input stimulus weighted by a spatial mask or ‘receptive field’ and sum the resulting values. This number, loosely analogous to a membrane potential, is then fed through a nonlinearity to approximate rectified neural firing rates. The model also included two sources of noise: (1) a pixel-wise input noise, which models stochasticity in phototransduction; and (2) an output noise, which models stochasticity in output firing rates. A key difference between the two is that the input noise is subject to the nonlinearity, and thus its effect becomes stimulus-dependent, while the output noise is not. We trained the model by optimizing mutual information under the constraint of a 1 activation (over all images) for each neuron (see Methods). This constraint is intended to model the metabolic cost of generating action potentials.

### Receptive field tiling is predicted by efficient coding

Following training, units in the model exhibited either ON-center or OFF-center receptive fields with strongly rectified nonlinearities (**Fig 1B-C)**. Each population of ON-center units and OFF-center units (referred henceforth as ON cells and OFF cells, respectively) exhibited a mosaic-like organization: their receptive fields tiled space (**Fig 1D)** Karklin and Simoncelli (2011). When varying noise levels, we reliably found circular ON and OFF receptive fields with input noise between 0.00 and 0.75 and output noise between 0.75 and 3.00. Input and output noise levels outside these ranges produced irregular receptive field shapes (**Sup Fig 1)**, as did nonnatural distributions of images (**Sup Fig 7)**. Therefore, we restricted the analysis to this range of noise regimes and focused on natural image patches.

### Mosaic arrangements depend on input and output noise

We next analyzed how these optimized mosaics of ON and OFF receptive fields were spatially arranged with respect to one another. Each optimized mosaic formed a nearly hexagonal grid (**Fig 1D)**. Analysis of the spatial relationships between the mosaics in **Fig 1D-E** revealed they were anti-aligned (**Fig 2D)**. However, as we explored the effect of changing the amounts of input and output noise on the optimization of this model, we observed other mosaic combinations that were aligned (e.g. **Fig 2D)**. Generally, small input and output noise values yielded aligned mosaics, while larger values produced anti-aligned mosaics (**Fig 2A-D)**. Importantly, for a fixed amount of input and output noise, multiple instances of the optimization produced consistent results, indicating that mosaic alignment and anti-alignment depended on the specific amount of input and output noise, and did not depend on different initializations to the optimization. Moreover, these same effects held when image patches were twice as large (**Sup Fig 8)**, arguing against simple edge effects. These results motivated a more thorough analysis of how the relative spatial organization between the two mosaics depended on input and output noise.

### A simplified ‘one-shape’ model captures the noise dependence of mosaic arrangements

To examine the dependence of the mosaic arrangements on noise over a range of densely sampled values, we utilized a simpler ‘one-shape’ model that trained much more quickly than the full model (see Methods). This model assumed that all receptive fields shared a common shape (a difference of concentric Gaussians), while optimizing over both the parameters of this shape and the locations of the receptive fields (see Methods). The learned kernels of this simplified model also exhibited ON and OFF receptive fields (**Sup Fig 2A)** and ON and OFF mosaics (**Sup Fig 2B)**. In addition, the optimal radial kernel exhibited a center-surround organization (**Sup Fig 2C)**. To further speed training, we also reduced the number of units in the model to 7 ON and 7 OFF cells (their polarities were fixed during the optimization). These simplifications allowed us to rapidly judge if the optimization preferred alignment or anti-alignment (**Fig 3)**. To ensure these simplifications did not strongly bias the results, we compared the mosaic arrangements for particular input and output noise combinations across the one-shape model and the full model. In all cases examined, the two models produced matching aligned or anti-aligned results (**Sup Fig 3)**. Thus, the one-shape model is a useful proxy for larger-scale and more general optimizations and could be used to rapidly and reliably determine if mosaics were aligned or anti-aligned for a given set of noise parameters.

### Mosaics exhibit a phase change as a function of input and output noise

To determine the optimal arrangement of ON and OFF mosaics for different amounts of input and output noise, we performed a grid search over input and output noise values, using the one-shape model. We ran the model with input noise ranging from 0.00 to 0.75 in steps of 0.05, and output noise from 0.75 to 3.00 in steps of 0.10. Optimizing the location of seven receptive field centers in a circular space nearly always resulted in a single kernel in the center surrounded by the remaining six kernels at the vertices of a hexagon. (see **Fig 3)**. When output noise level was low, ON and OFF receptive field mosaics were aligned as in the first row of **Fig 3B** and resulted in near-zero median nearest neighbor distances (dark purple colored area in **Fig 3A)**, whereas ON and OFF mosaics were anti-aligned under higher output noise levels as in the last row of **Fig 3B** and resulted in larger median nearest neighbor distances (lighter colored area in **Fig 3A)**. In particular, this analysis reveals an abrupt transition between mosaic alignment and anti-alignment (**Fig 3A)**. Thus, our model optimization indicates the solution to the sensor alignment problem depends on noise present in the encoding process. Below, we build an analytical model for understanding this result.

### Retinal phase transitions from an analytic model of efficient coding

To understand the factors that give rise to the phase transition between aligned and anti-aligned mosaics, we developed a simplified 1-dimensional analytic model in which ON and OFF grids have identical nonlinearities, fixed kernels, fixed spacing, and are only allowed to shift relative to one another (see supplementary section 2). Moreover, we assume that images are drawn from a Gaussian distribution with a 1/*f* frequency spectrum that is the 1D equivalent of the two-dimensional scale-free distribution of natural images Atick and Redlich (1990), Ruderman (1994), though other frequency distributions produced the same effect (**Sup Fig 9)**. We verified via simulation that, even with all of these restrictions, the phase transition occurs just as in the less constrained models in both one and two dimensions, suggesting that the remaining free parameters — gain, threshold, and mosaic alignment — are sufficient to account for the phase transition (**Sup Fig 5** and **Sup Fig 6)**.

To gain additional insight into the origins of the transition, we analyzed the behavior of this model in the more analytically tractable limit of near-independent neurons. The starting point for this analysis is a recognition that the conditional entropy between stimulus and neural response can be conveniently written as a sum of single-neuron terms and higher-order corrections:

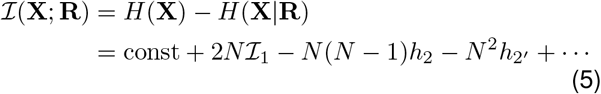

Here, *H*(*X*) is the (constant) entropy of the stimulus, 2*N* is the number of ON + OFF cells, 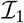 is the contribution of each neuron individually, and *h*_2_ and *h*_2′_ are correction terms due to non-independence (and thus redundancy) of ON-ON/OFF-OFF (same polarity) and ON-OFF (opposite polarity) cells, respectively (see supplementary section 2B). Importantly, these corrections are always negative and vanish for widely separated cells. For our analysis, we focus only on these leading order terms, corresponding to an assumption of approximately independent neurons with small pairwise interactions. While this assumption is violated for natural images, which possess long-range correlations and induce triplet and higher-order interactions, our analytical results are nonetheless matched by simulations in the full model, suggesting that intuitions gleaned from this approximation are sufficient to explain the transition between alignment and anti-alignment. In the following two sections we show (1) that noise and image statistics set the optimal neuron response thresholds, and (2) that these thresholds control the amount of redundancy in the population responses, thereby dictating whether mosaic alignment or anti-alignment is the preferred state.

### Output noise and image outliers drive increases in optimal neuron response thresholds

First, we considered optimizing only the 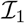 term above, which captures the contribution of each neuron individually to encoding stimulus information. In particular, we focused on the influence of output noise and the distribution of natural images on the output nonlinearities of the neurons (we focused on output noise because the transition depended much more on output than input noise, see **Fig 3A)**:

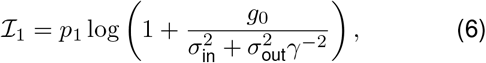

where *g*_0_ is a constant that depends on the image distribution and the receptive field shape, *p*_1_ is the probability that the neuron is active, *γ* is the neuron’s gain, and these latter two are related through the constraint that the average response of the neuron is 1 (see supplementary section 2B, Equation 18). As the output noise of neurons increases, maximizing 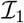 dictates that neurons should respond by increasing their firing threshold (**Fig 4A)**. This allows the neurons to reserve spikes for stimuli with high signal-to-noise ratios Laughlin (2001). Thus, units respond less often but do so with higher gain when active, thereby mitigating the impact of noise. In addition, the optimal firing threshold depends on the nature of the image distribution. For neuron activation distributions with more outliers (see Methods), thresholds increase as the tail of the distribution becomes heavier (**Fig 4B)**. Intuitively, this is because heavier-tailed distributions require higher thresholds to maintain the same overall response probability. Finally, it can be shown that decreasing the input noise has the effect of increasing the effective threshold (see supplementary section 2B, Equation 19). Thus, the optimal spiking threshold is determined both by the input and output noise and the distribution of natural scenes (schematized in **Fig 4C)** Gjorgjieva *et al.* (2014).

**FIG. 4.**
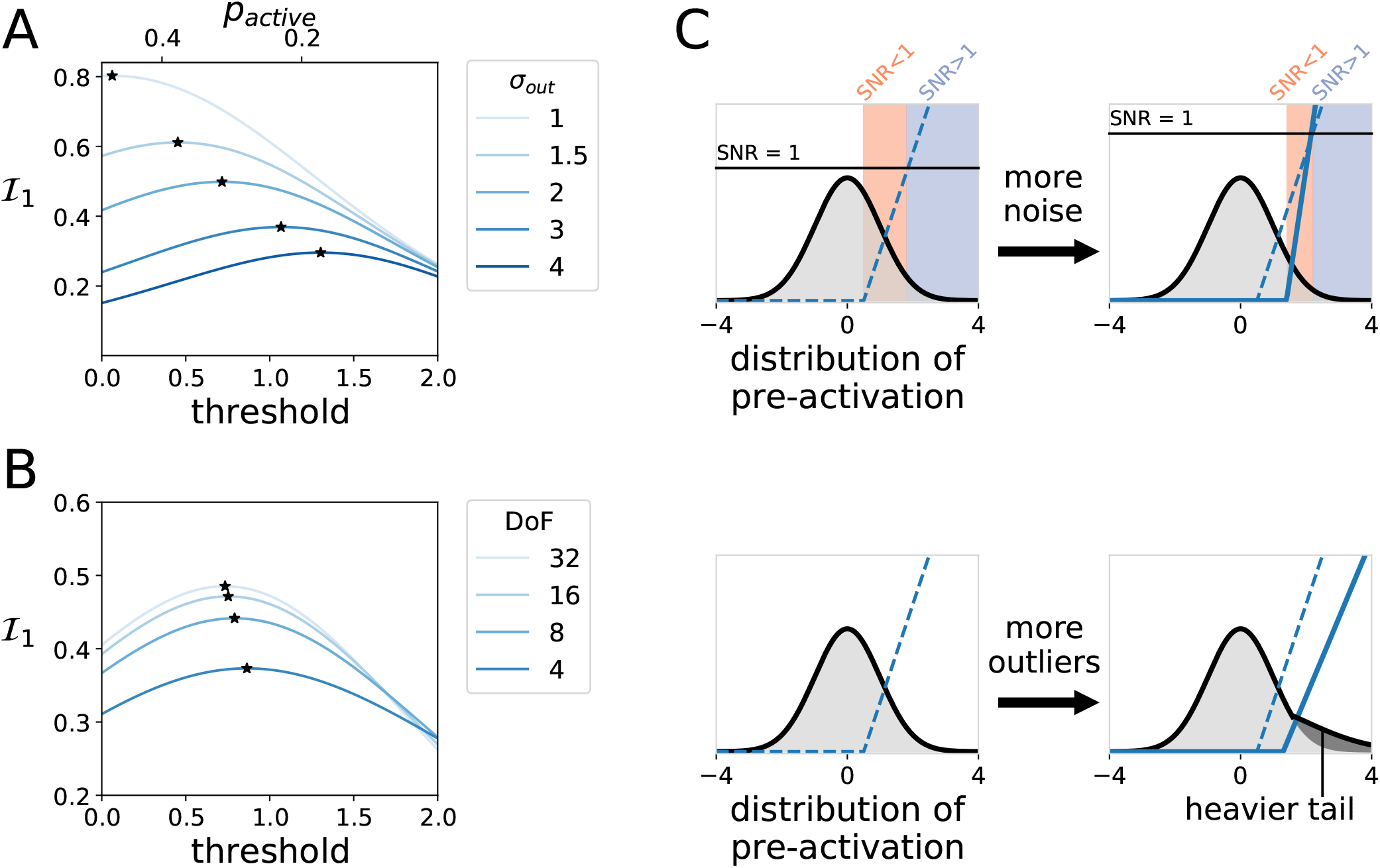
Optimal threshold increases with higher noise and heavier tailed image distributions. **A-B.** Contribution of a single neuron to the mutual information as a function of threshold. Stars mark optimal thresholds for multiple parameter values. **A.** When the input distribution is Gaussian, the optimal threshold increases with higher output noise, even as overall information decreases. **B.** When the input distribution is heavier tailed, modeled with Student’s t distributions with varying degrees of freedom, the optimal threshold again increases. **C.** Schematic illustrating the effects of increased noise or outliers. With increased output noise, neurons’ signal-to-noise (SNR, black line, upper right) decreases, and efficient coding predicts that units should increase threshold and gain to reduce low SNR responses (peach). Similarly, when pre-activations are heavy-tailed (lower-right), efficient coding predicts that thresholds should increase and gains slightly decrease (see **Sup Fig 11)**, since more mass is contained in outliers. Thus, both output noise and heavy tails lead to higher thresholds.

### Mosaic anti-alignment achieves redundancy reduction at high neuron thresholds

Next, we considered the effect of ON-OFF pair corrections to Equation 5. These terms, represented by – *N*^2^*h*_2′_ above, capture the effects of redundancy in ON-OFF encoding: they are always negative and are the only contributions to mutual information that change as a function of alignment. Thus, while the overall trend is that mutual information decreases with increasing threshold, mosaic configurations may be more or less efficient for information transmission as they limit the impact of these terms. More specifically,

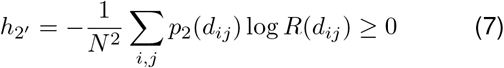

Where *d_ij_* is the distance between ON cell *i* and OFF cell *j, p*_2_ is the probability that the pair is coactive, and 0 < *R* < 1 is a function that is approximately independent of both output noise and neuron nonlinearity (see supplementary section 2B). This correction term is small for large separations, large around the intra-mosaic spacing, and almost zero in a region around zero inter-cell distance (**Fig 5A)**. Importantly, the width of this ‘independence zone’ around zero inter-cell distance widens as output noise grows larger. This is explained by the fact that, as output noise increases, so does the threshold (**Fig 4A)**, and at large threshold values, nearby opposite polarity pairs are almost never coactive (see supplementary section 2B).

**FIG. 5.**
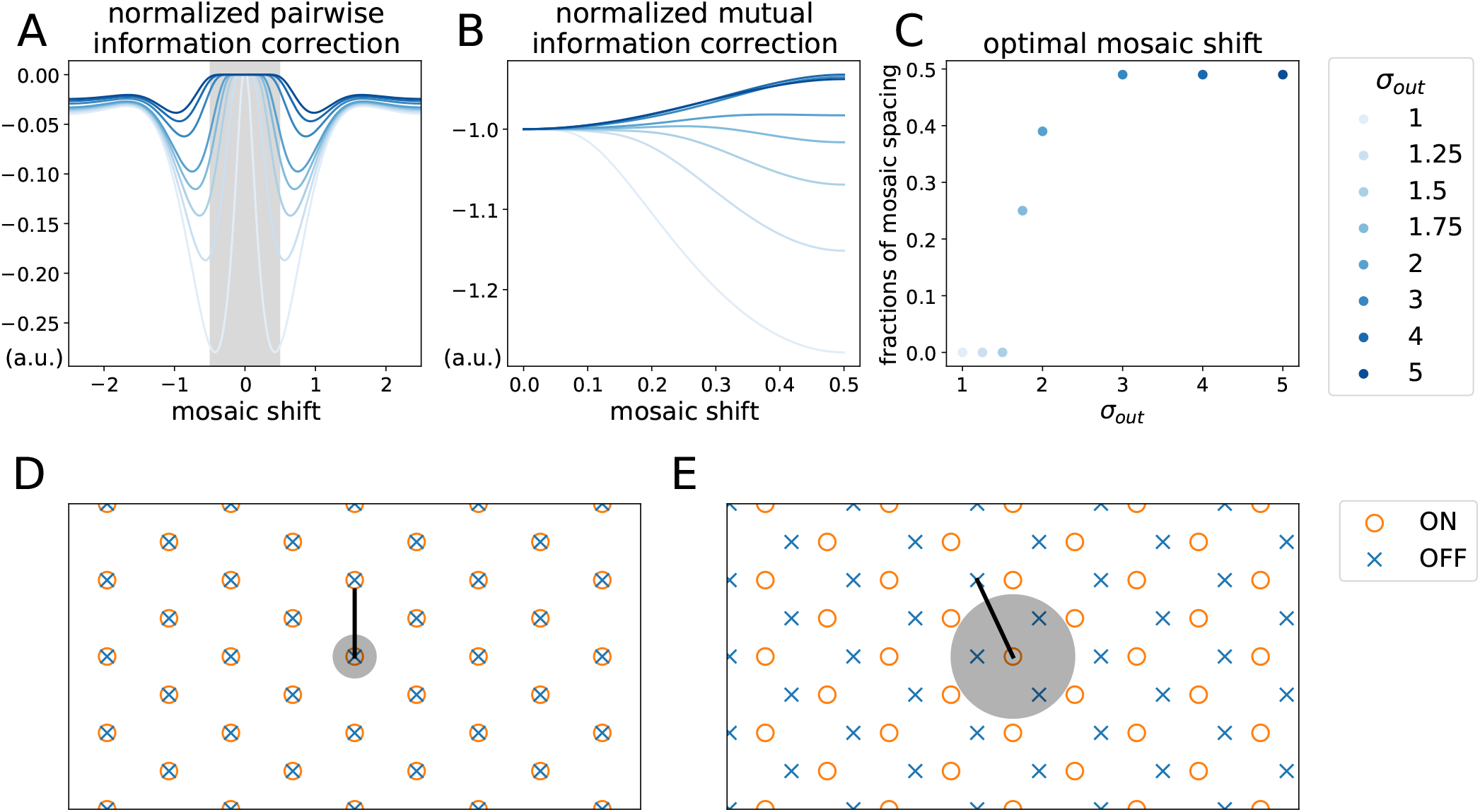
Pairwise mutual information corrections explain the phase transition between alignment and anti-alignment. **A.** Normalized pairwise information correction 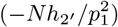 for pairs of ON and OFF neurons as a function of mosaic shift. One unit is the spacing between nearest-neighbor receptive field centers of the same polarity. With low output noise, an aligned position (i.e., no shift) is the only placement to avoid a large negative information correction, while with higher output noise, an ON-OFF pair can still have a finite shift within the gray area without incurring any penalty. **B.** The total normalized pairwise information correction as a function of mosaic shift, assuming equispaced ON and OFF receptive field centers filling the 1-D space. With low output noise, any mosaic shift lowers mutual information, while with high output noise, the pairwise mutual information becomes less negative when the mosaic is shifted by 0.5, i.e. when anti-aligned. **C.** The optimal mosaic shift as a function of output noise. For *σ*_out_ = 1.7, *σ*_out_ = 2, the optimal mosaic shift lies between 0 (alignment) and 0.5 (anti-alignment), resembling the arrangements we obtained in the middle row of **Fig 3B. D.** Schematic of optimal 2-D mosaics in a low-noise regime. The gray circle indicates the area within which an OFF cell can be located with zero pairwise correction to mutual information. **E.** Schematic of optimal 2-D mosaics in a high-noise regime. Here, the gray area has grown (as in **A**), allowing three neighbors of the opposite polarity to fit inside without loss in mutual information.

This analysis gives rise to the following account of the phase transition: at low levels of output noise or light-tailed distributions of neuron activations, neuronal thresholds are low, and only aligned receptive fields avoid the large *N*^2^*h*_2′_ penalty. Thus, in one dimension, each ON cell is non-redundant (save one bit of sign information; supplementary section 2C) with its aligned OFF partner, but image correlations make it highly redundant with the two OFF (and two ON) cells on either side. Likewise, in two dimensions, each ON cell is non-redundant (save one bit of sign information; see supplementary section 2C) with its aligned OFF partner, but it suffers large negative corrections to encoding efficiency from its six other OFF neighbors. However, as activity thresholds rise, a finite-sized region develops around each cell within which other receptive fields of opposite polarity are almost never coactive (**Fig 5A**, gray). This region of approximate independence grows with threshold, to the point that, when it is sufficiently large, more than one OFF cell can fit inside when the mosaics are antialigned (**Fig 5D-E, gray circle**). Thus, anti-alignment results in lower redundancy (via reduced *h*_2_ penalties) and increased mutual information. This picture is born out quantitatively by the model: as output noise (and thus threshold) increases, the optimal shift interpolates smoothly between aligned and anti-aligned configurations (**Fig 5B-C** and cf. **Fig 3B**, second row), suggesting a second-order Landau-Ginzburg phase transition Binney *et al.* (1992).

### Outliers in natural image statistics modify the transition from aligned to anti-aligned mosaics

The analysis above identifies output noise and image (stimulus) outliers as driving optimal encoding strategies toward higher neural response thresholds, which we claim is the key factor mediating the phase transition from aligned to anti-aligned mosaics. However, this analysis makes numerous simplifying assumptions, including a restriction to one dimension. Thus, we sought to test whether these predictions generalized to a less-restricted two-dimensional model in which image statistics were manipulated.

First, we identified image patches that generated the highest and the lowest firing rates in our fitted model (**Sup Fig 4)**. We identified these images using the firing rates under the two noise regimes, one that produced aligned and one that produced anti-aligned mosaics (**Fig 2B** (*σ*_in_ = 0.1, *σ*_out_ = 1.0) and **Fig 2D**(*σ*_in_ = 0.4, *σ*_out_ = 3.0)). In both noise regimes, the 100 image patches producing the highest firing rates were extreme luminance values (nearly all black or all white) or extreme contrast values (e.g. contained one or more edges between black and white regions). On the other hand, the 100 images producing the lowest firing rates were nearly homogeneous gray patches (**Sup Fig 4)**. A 2-D histogram of the mean and standard deviation across the pixels of all of the available sample image patches confirmed that these images exhibited outlier mean or contrast values (**Sup Fig 4E)**. Thus, under both noise regimes, the largest firing rates (across all neurons) were produced by rare, outlier images.

We next tested what impact these outlier images exert on the phase transition between aligned and anti-aligned mosaics. We re-ran the ‘one-shape’ model (e.g. **Fig 3A)**, but with the altered image sets that either reduced or increased the frequency of outlier images. From the distribution of all of the 18,587,848 possible image patches in the dataset (**Fig 6A)**, we considered a 2-D space composed of the z scores of the patch mean and SD values, and we drew the unit circle in this 2-D space. Within this space, we considered regions within 1 to 3 standard deviations from the mean of the average pixel intensity (per image) and from the mean of the variance over pixel intensities (per image). We then trained the one-shape model (see **Fig 3)** on only the patches within each boundary (**Fig 6B)**. As predicted, we found that the transition between aligned and anti-aligned mosaics occurred at lower noise levels as more outlier images were included in the training set and occurred at higher noise levels when these outlier images were removed. Furthermore, these changes were mediated by an increase in activation threshold as outliers grew more prevalent (**Fig 6C)**. Thus, we were able to reproduce the effects of outlier images on the phase transition between aligned and anti-aligned mosaics.

**FIG. 6.**
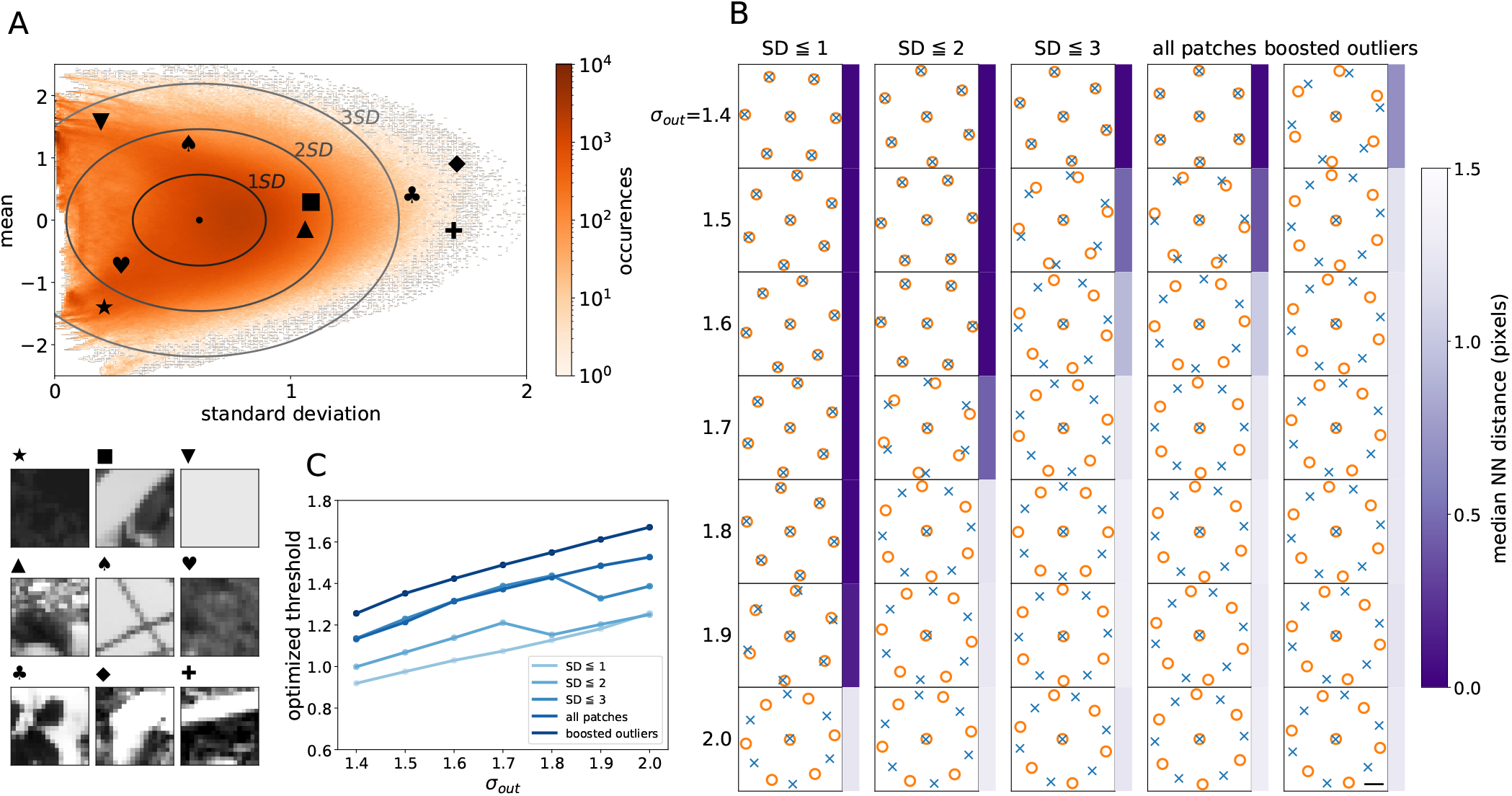
Outlier image patches drive an anti-aligned relationship between ON/OFF receptive field mosaics via higher optimal thresholds. **A.** Distribution of all 18,587,848 image patches from the dataset, plotted as a 2D-histogram with respect to the mean and the standard deviation of each image patch (orange). A few example patches are shown in the lower left, with the corresponding locations in the histogram marked with matching symbols. The means are centered at zero (sd =0.575), and the standard deviation values are centered at 0.623 (sd = 0.187). Ellipses in the histogram represent data within 1-3 standard deviations of the mean, which we use to construct artificial datasets with varying contributions of outliers. **B.** Optimized ON and OFF receptive field mosaics using the image patches within the boundaries denoted in **A** with *σ*_in_=0.4, *σ*_out_=1.4 to 2.0, plotted with the same scheme as in **Fig 3.** As predicted, when trained on image sets that systematically excluded outliers (i.e., had lighter-tailed distributions), phase transitions happened only at larger values of output noise. The fourth column denoted as “all patches” used the full dataset as in **Fig 3.** The last column, denoted as “boosted outliers,” used an augmented dataset to which we added horizontal and vertical mirror flips of patches outside the 1.5 SD circle, thereby creating a distribution with even more outliers. This corresponds to the top 20% of z-scores among all patches in the dataset. Here, the phase transition happened at lower output noise values. The scale bar shows a 1-pixel distance. **C.** Optimal thresholds as a function of output noise, using input distributions with different levels of outliers. As predicted, the optimal thresholds tended to increase as the input distribution contained more outliers, causing the phase transition to happen at a lower output noise level.

## DISCUSSION

Efficient coding theory provides a basis for understanding many features of early sensory processing. This includes the structure of individual receptive fields, how receptive fields adapt, and their mosaic-like arrangement to uniformly tile space Barlow (1961), Doi *et al.* (2012), Atick and Redlich (1992), Karklin and Simoncelli (2011), Ratliff *et al.* (2009). We have extended this framework to investigate how the receptive fields of distinct cell types should be spatially arranged, given sensory noise and the statistics of natural stimuli. We term this the ‘sensor alignment problem.’ Below, we summarize the features and properties of the model that produce center-surround receptive fields, mosaics, and mosaic coordination.

### a. Center-surround receptive fields

The convergence of the filters to center-surround receptive fields was robust over a range of input and output noise values (**Sup Fig 1)**. However, perhaps surprisingly, convergence to center-surround filters required the higher-order correlations present in natural scenes: training on Gaussian images with the same covariance as the natural scenes did not yield center-surround filters (**Sup Fig 7)**.

### b. Mosaics

In all cases where the model produced center-surround receptive fields, it likewise produced ON and OFF cells and mosaics (see above). Moreover, when center-surround receptive fields were enforced by parameterizing filters as a difference of Gaussians, mosaics always resulted (**Sup Fig 2, 3, 5, 9)** even without higher-order or even long-range correlations (**Sup Fig 9)**.

### c. Mosaic coordination

The phase transition between aligned and anti-aligned mosaics occurred under a wide range of conditions, including natural and synthetic image sets. Gaussian-distributed images were sufficient to drive the phase transition between the aligned and anti-aligned states (**Sup Fig 5B**). Indeed, this remained true even for images with only short-range correlations (**Sup Fig 9)**, albeit only at extremely high levels of noise.

From these relationships, we can sketch the following qualitative account of the phase transition between aligned and anti-aligned mosaics: Given center-surround receptive fields, efficient coding argues that neurons should maximize information transmission by both distributing receptive fields to encode as much unique information as possible and reducing redundancy, which occurs when nearby cells are coactive to the same stimulus. The first intuition leads to the formation of mosaics, while the second leads, at low noise levels, to mosaic alignment. This is because optimal neuron response thresholds in the low-noise regime are lower (**Fig 4A)**, so nearby ON and OFF cells are likely to be coactive unless they are aligned, and this redundancy cost outweighs the benefits of encoding slightly different locations in the visual field (**Fig 5D)**.

However, in the high-noise or heavy-tailed neural response regimes, optimal encoding requires that individual neurons raise their response thresholds (**Fig 4A)**. This results in the decorrelation of nearby cells Pitkow and Meister (2012), creating an ‘independence zone’ around each cell that grows with noise (**Fig 5A, E)**. Receptive fields located within this distance of one another are nearly independent in their responses, allowing for nearby cells to sample distinct spatial locations without reducing encoded information through redundancy (**Fig 5A)**. The optimal configuration in this regime then becomes mosaic anti-alignment across a wide variety of conditions (**Fig 6, Sup Fig 9)**.

In previous work we have shown that real retinal mosaics of ON and OFF cells that encode similar visual features are anti-aligned Roy *et al.* (2021). Here, we have shown that the optimal solution to the sensor alignment problem depends on both system noise and stimulus statistics. Thus, the anti-alignment of retinal mosaics suggests that retinal processing is optimized for low signal-to-noise conditions such as detecting dim or low contrast stimuli Dhingra *et al.* (2003), Field and Sampath (2017). In natural environments, the retina is probably faced with both high and low SNR conditions. When SNR is high, suboptimal processing that squanders some signal probably has little impact on sensory encoding because signal is plentiful. However, when SNR is low, the signal is precious and optimal processing is required to spare the signal from noise. This raises the question: How much does anti-alignment improve encoding under low SNR conditions? We previously found an ≈ 4% increase in mutual information for antialigned mosaics over aligned mosaics Roy *et al.* (2021). While small, it is worth noting that the presence of a surround – a feature that is thought to be very important for optimal encoding of natural scenes – only improves encoding by ≈ 20%. Thus anti-alignment of mosaics is roughly ≈ 20% as important as center-surround receptive field structure.

This study connects to several other strands of work on efficient coding in the retina. As noted above, efficient coding as redundancy reduction subject to signaling costs gave theoretical weight to the idea of center-surround receptive fields as whitening filters for visual stimuli Barlow (1961), Atick and Redlich (1990, 1992), Laughlin (2001), Srinivasan *et al.* (1982) and are thus matched to the statistics of natural scenes. However, as later work has made clear, nonlinearities, particularly response thresholds, are perhaps even more important in optimizing retinal codes Gjorgjieva *et al.* (2014), Pitkow and Meister (2012), Gjorgjieva *et al.* (2019). In addition, noise in both sensory transduction (inputs) and neural outputs affects coding efficiency Atick and Redlich (1990, 1992), Röth *et al.* (2020). This work, by placing all of these factors in an optimization-based framework Karklin and Simoncelli (2011), allows for investigating the relative importance of each factor in determining the optimal spatial layout of receptive fields. Importantly, this approach relies only on information maximization arguments, and the optimization does not rely on fitting data from electrophysiological recordings, as required by recent deep learning models of retinal coding Maheswaranathan *et al.* (2018), Ocko *et al.* (2018).

By focusing on the factors that govern the transition between aligned and anti-aligned mosaics from an optimal encoding standpoint, we have necessarily ignored several real biological complexities. First and foremost, the retina contains roughly 40 distinct RGC types. Our current efficient coding model only generates two types of units that encode similar visual features, but with opposite polarity, and thus we are not yet able to determine whether RGC types that encode distinct visual features should be independent or coordinated. But a limited inspection of the mosaic relationships across RGC types that encode distinct visual features suggested that the measured receptive field mosaics were statistically independent Roy *et al.* (2021). Developing an efficient coding model that is optimized to encode natural movies may yield a greater diversity of cell types Ocko *et al.* (2018). Many RGC types that encode particular spacetime features such as direction of motion or ’looming’ detectors also form mosaics Vaney *et al.* (2012). Whether these detector grids are coordinated or should be coordinated remains an open question. Moreover, our analysis does not consider constraints on development that result in irregular mosaics, which have been shown to require local changes in receptive field shape for optimal encoding Liu *et al.* (2009). Nonetheless, during the optimization process, we do observe such local changes as receptive fields push against one another during mosaic formation. Finally, our results motivate understanding the implications of efficient coding theory to other other sensory systems such as touch receptors on the skin: might they be spatially coordinated for the efficient coding of touch Kuehn *et al.* (2019)? More generally, these observations demonstrate that efficient coding theory can make predictions about emergent properties present in the organization of the nervous system, such as how large populations composed of multiple cell types should be arranged to optimally encode the natural environment.

## ACKNOWLEDGMENTS

We thank Joel Zylberberg, Fred Rieke, Nicolas Brunel, and Suva Roy for comments on the manuscript and helpful discussions. This work was funded by the National Institutes of Health, National Eye Institute grant R01 EY031396.

## I. SUPPLEMENTARY FIGURES

**Supplementary Figure 1.**
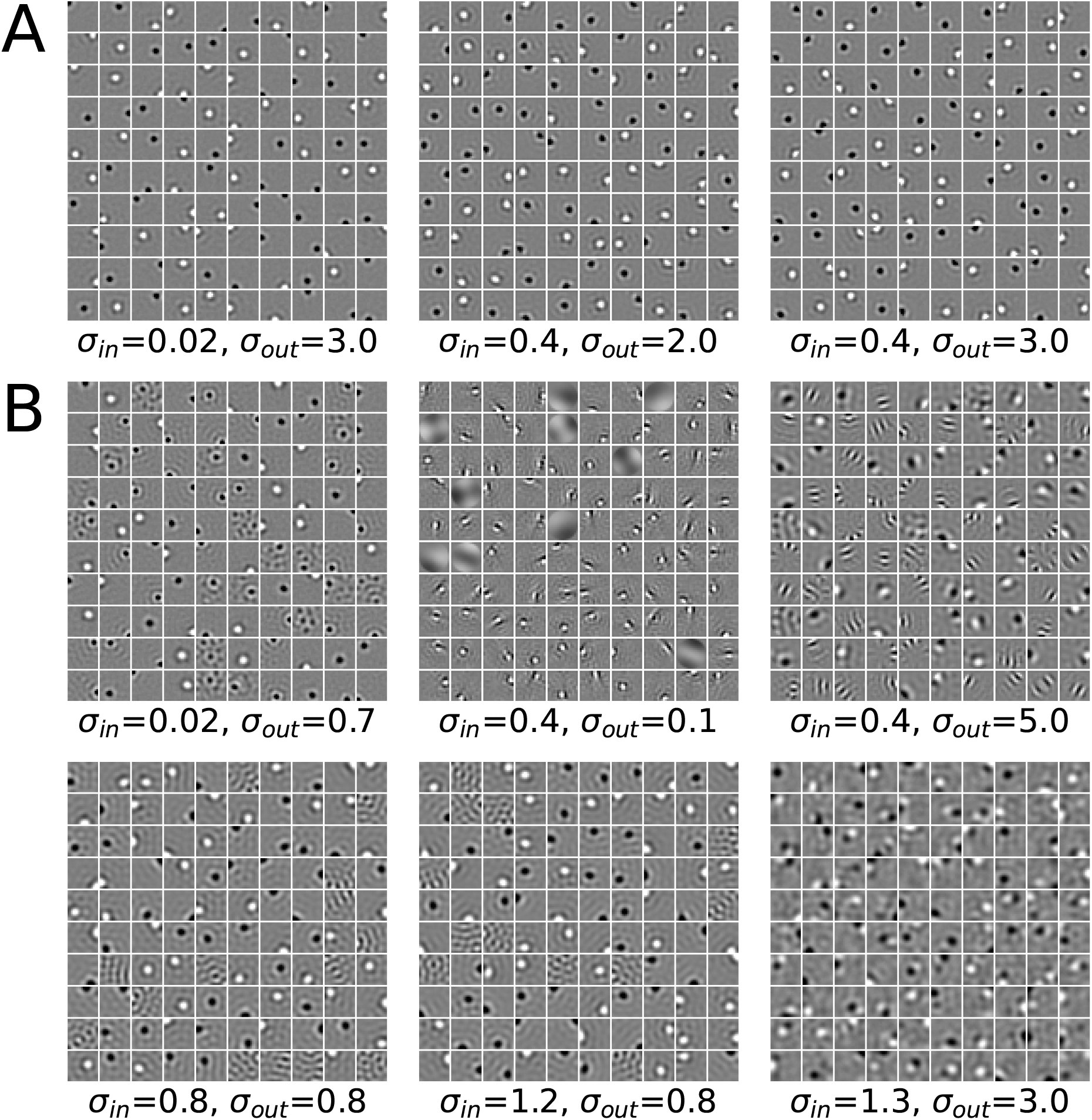
Circular ON and OFF receptive fields are robustly obtained within a range of input and output noise. **A.** With input noise in the range between 0.00 to 0.75 and output noise between 0.75 to 3.00, circular center-surround ON and OFF receptive fields are obtained reliably. **B.** Outside these ranges, i.e., when input noise is higher than 0.75 or output noise is lower than 0.75, either Gabor-shaped or noisy kernels are obtained.

**Supplementary Figure 2.**
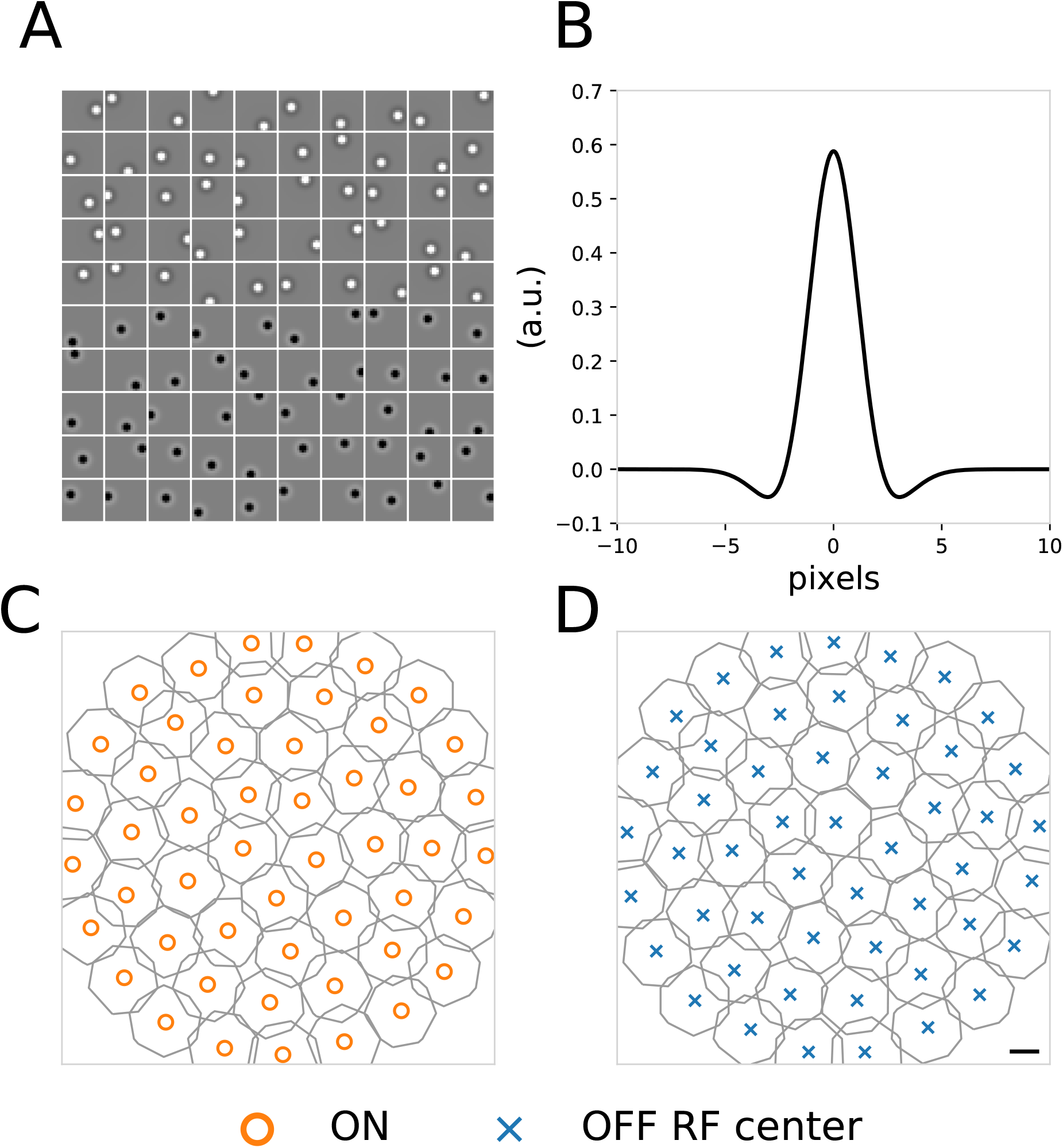
When all filters share the same kernel, filters form mosaics. **A.** A plot of 100 kernels that maximize information when all kernels are constrained to take the same shape but are free to move their centers (cf. **Fig 1C) B.** The radial shape function is parameterized as a difference of Gaussian curves. Learned radial shape parameters are: *a*=0.3332, *b*=0.1559, *c*=0.4121. **C.** Learned ON kernels sharing the same radial shape fill the field (similar to **Fig 1D)**. **D.** Learned OFF kernels sharing the same radial shape fill the field (similar to **Fig 1E)**. Scale bar is width of one image pixel.

**Supplementary Figure 3.**
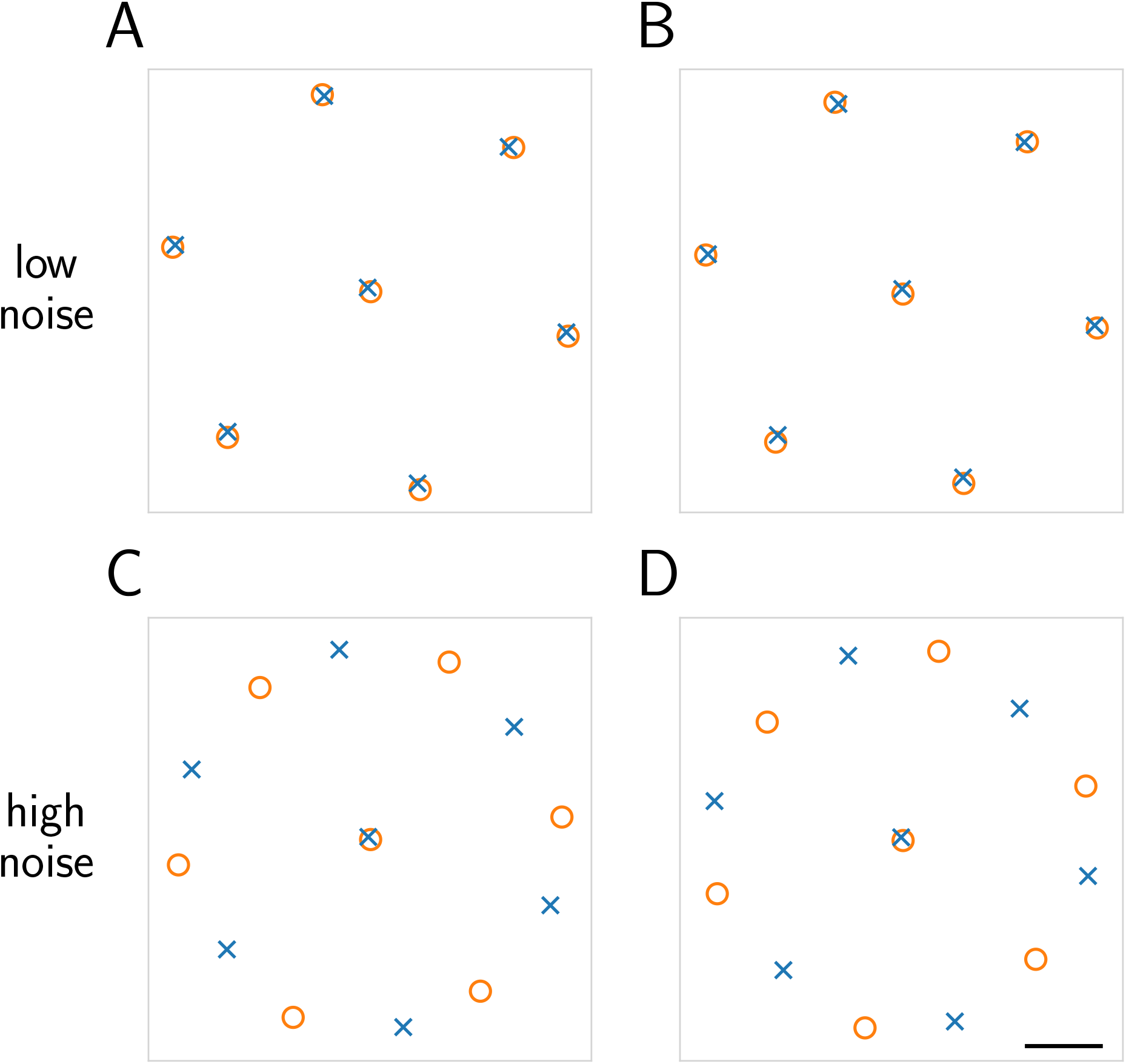
A one-shape model captures the dependence of mosaic arrangements on input and output noise. Receptive field centers for ON (orange circle) and OFF (blue x) cells under different sets of noise parameters. **A:** (*σ*_in_=0.02, *σ*_out_=1.0), **B:** (*σ*_in_=0.1, *σ*_out_=1.0), **C:** (*σ*_in_=0.4, *σ*_out_=2.0), **D:** (*σ*_in_=0.4, *σ*_out_=3.0). The first two parameter sets result in aligned mosaics, while the latter two, at higher levels of output noise, are anti-aligned. Scale bar is width of one image pixel.

**Supplementary Figure 4.**
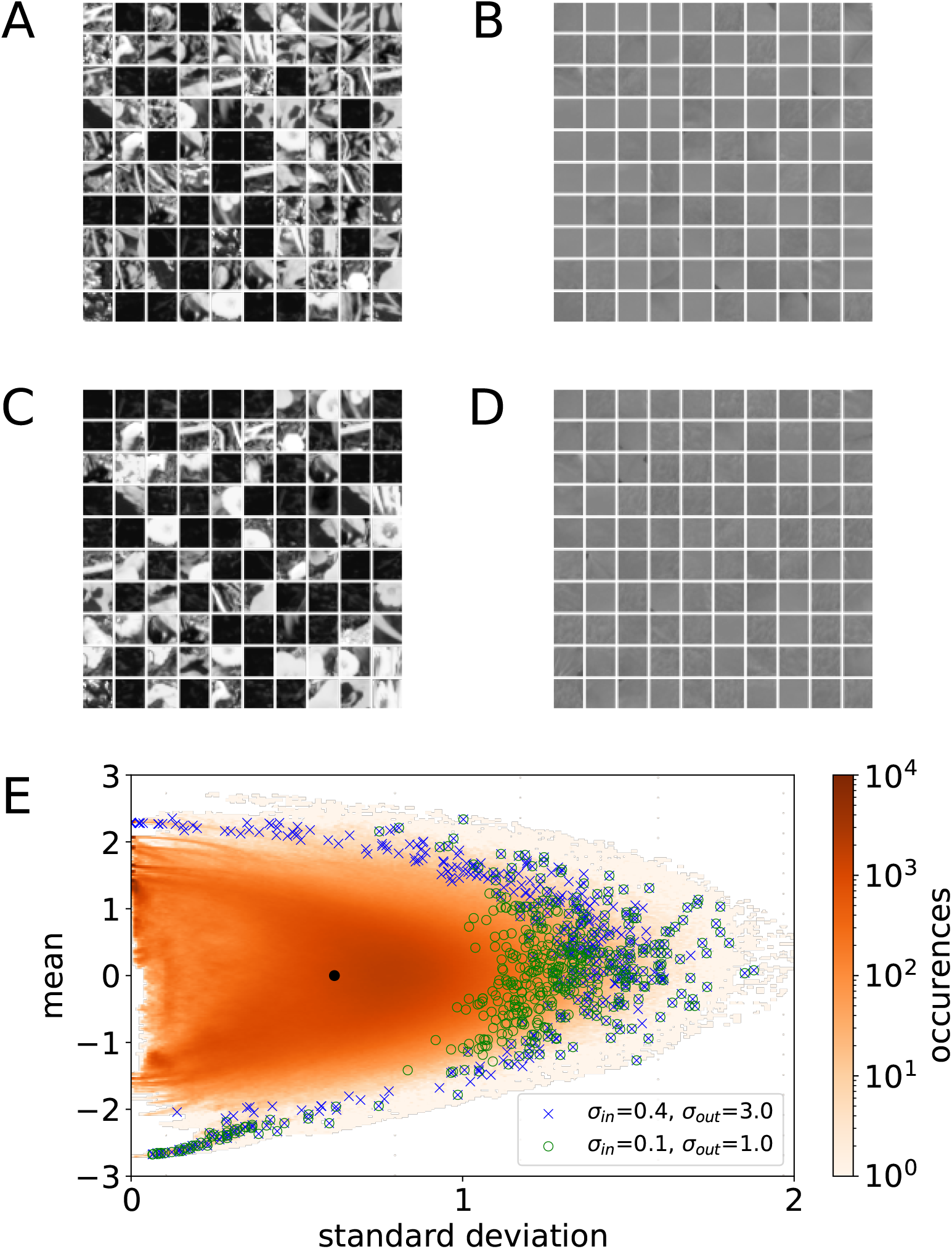
Image patches with the highest firing rates and lowest firing rates (among 100,000 randomly sampled image patches) are distinct in both the high noise (*σ*_in_=0.4, *σ*_out_=3.0) and the low noise (*σ*_in_=0.1, *σ*_out_=1.0) regimes. Most of the image patches with the highest firing rates are outliers of the overall image patch distribution, containing high contrast edges or particularly high or low mean intensity. This is also evident in firing rate outputs: Using the kernels optimized under the two noise regimes, as shown in **Fig 2B** and **Fig 2D**, we calculated the firing rates of 100,000 randomly sampled image patches and overlaid the top 100 patches with the highest firing rates over the z-score histogram in **Fig 6A. A.** 100 image patches with the highest firing rates under *σ*_in_=0.1, *σ*_out_=1.0. Images are either extremely dark or contain distinct edges, including those with more complicated edges compared to **C**, and had firing rates of 3.07±0.23. **B.** 100 image patches with the lowest firing rates under *σ*_in_=0.1, *σ*_out_=1.0. The 100 images with the lowest firing rates were also visually similar and had firing rates of 0.161±0.003. **C.** 100 image patches with the highest firing rates under *σ*_in_=0.4, *σ*_out_=3.0. Images are either dark or contain distinct edges, with firing rates of 4.39±0.54. **D.** 100 image patches with the lowest firing rates under *σ*_in_=0.4, *σ*_out_=3.0. Patches were almost entirely gray, with firing rates of 0.036±0.001. **E.** A 2-D histogram showing the mean and standard deviation of all of the available sample image patches (in orange, occurrences in log scale, same as **Fig 6A)** with the image patches in **A** (blue x) and the image patches in **B** (green o). In the low-noise condition, the outliers are a greater portion of the tail (green o).

**Supplementary Figure 5.**
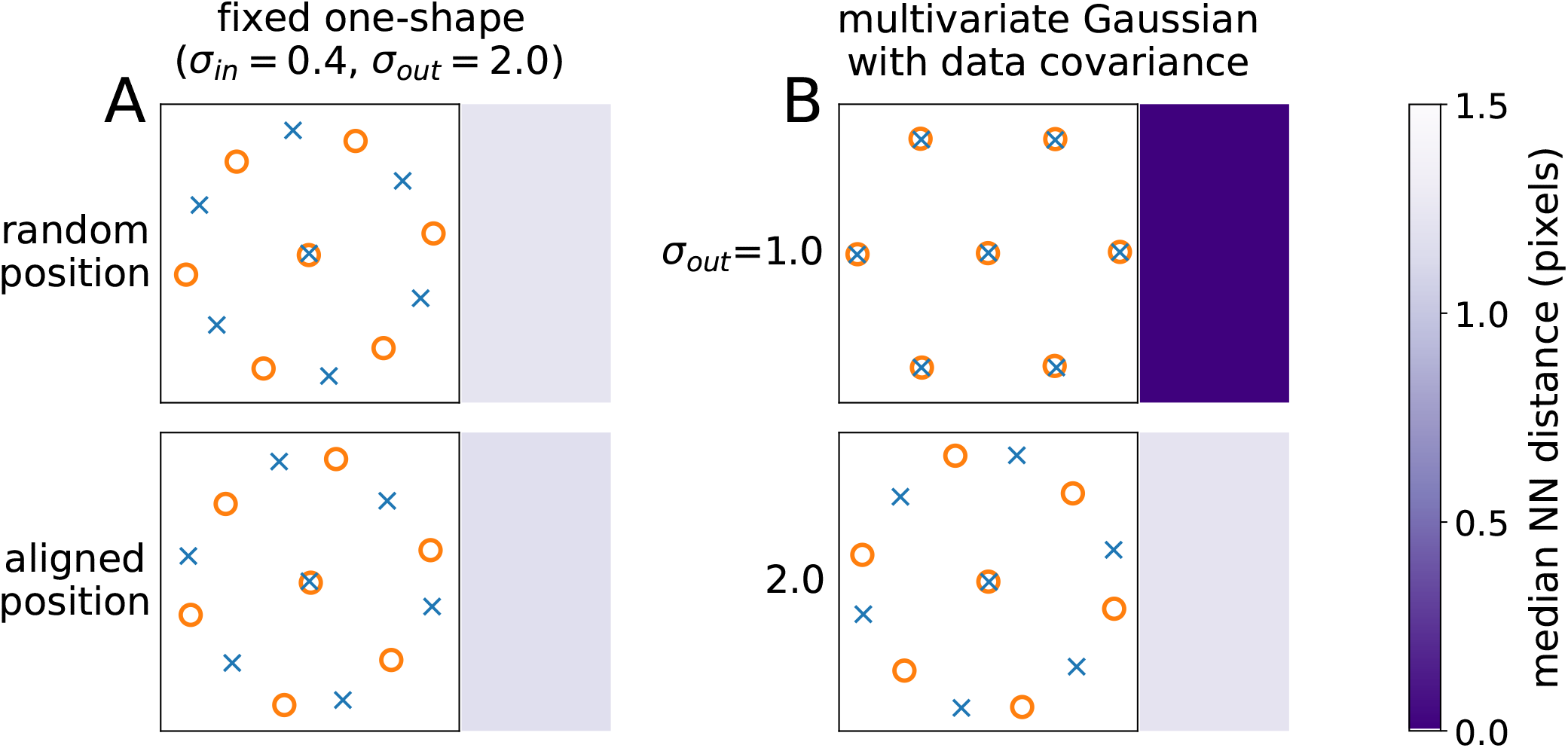
The transition between aligned and anti-aligned occurs when kernel shape is fixed and with Gaussian images. **A.** When kernel parameters are not learnable (fixed to *a, b, c* = [0.258, 0.162, 0.648]) and noise is high, the optimal configuration remains anti-alignment. This occurs when the receptive fields are initialized to random positions (top) or aligned (bottom) positions (cf. **Fig 5B** top row of “all patches”). Here *σ*_in_=0.4, *σ*_out_=1.0. **B.** The transition from aligned to anti-aligned mosaics occurs when images are drawn from a covariance-matched Gaussian distribution. This matches the assumptions of the analytical model (see Appendix) and demonstrates that heaver-than-Gaussian tails are not necessary for anti-alignment to be the favored state.

**Supplementary Figure 6.**
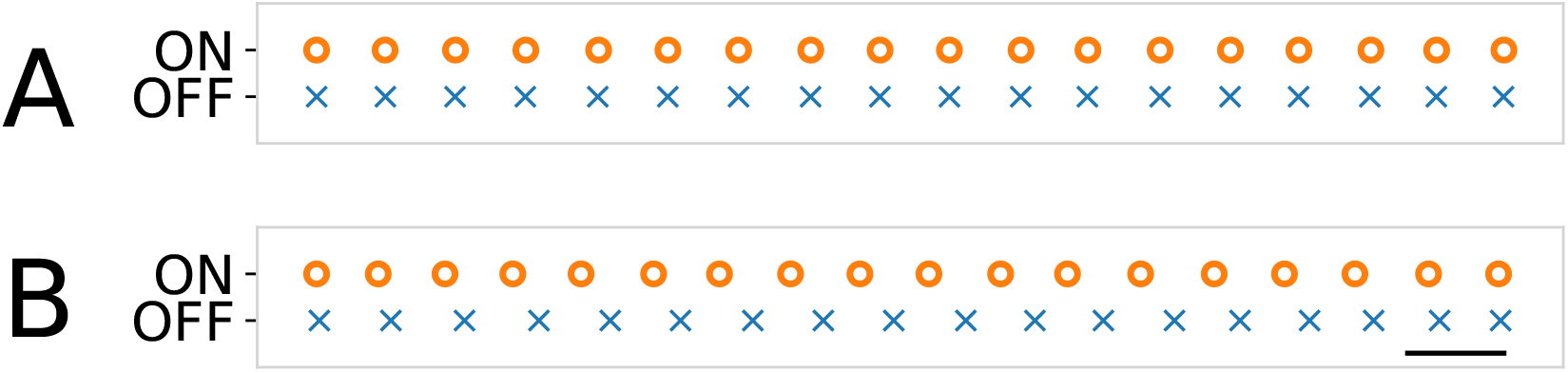
The phase transition occurs in a 1-dimensional model. Optimal receptive field centers of 18 ON and 18 OFF receptive fields in a 1-by-64 pixel input. Receptive field centers for ON (orange circle) and OFF (blue x) cells are optimized under two sets of noise parameters, A: (*σ*_in_=0.4, *σ*_out_=1.0), B: (*σ*_in_=0.4, *σ*_out_=3.0). Noise parameters in **A** produce aligned mosaics, while those in **B** produce anti-aligned mosaics. Scale bar is width of one image pixel.

**Supplementary Figure 7.**
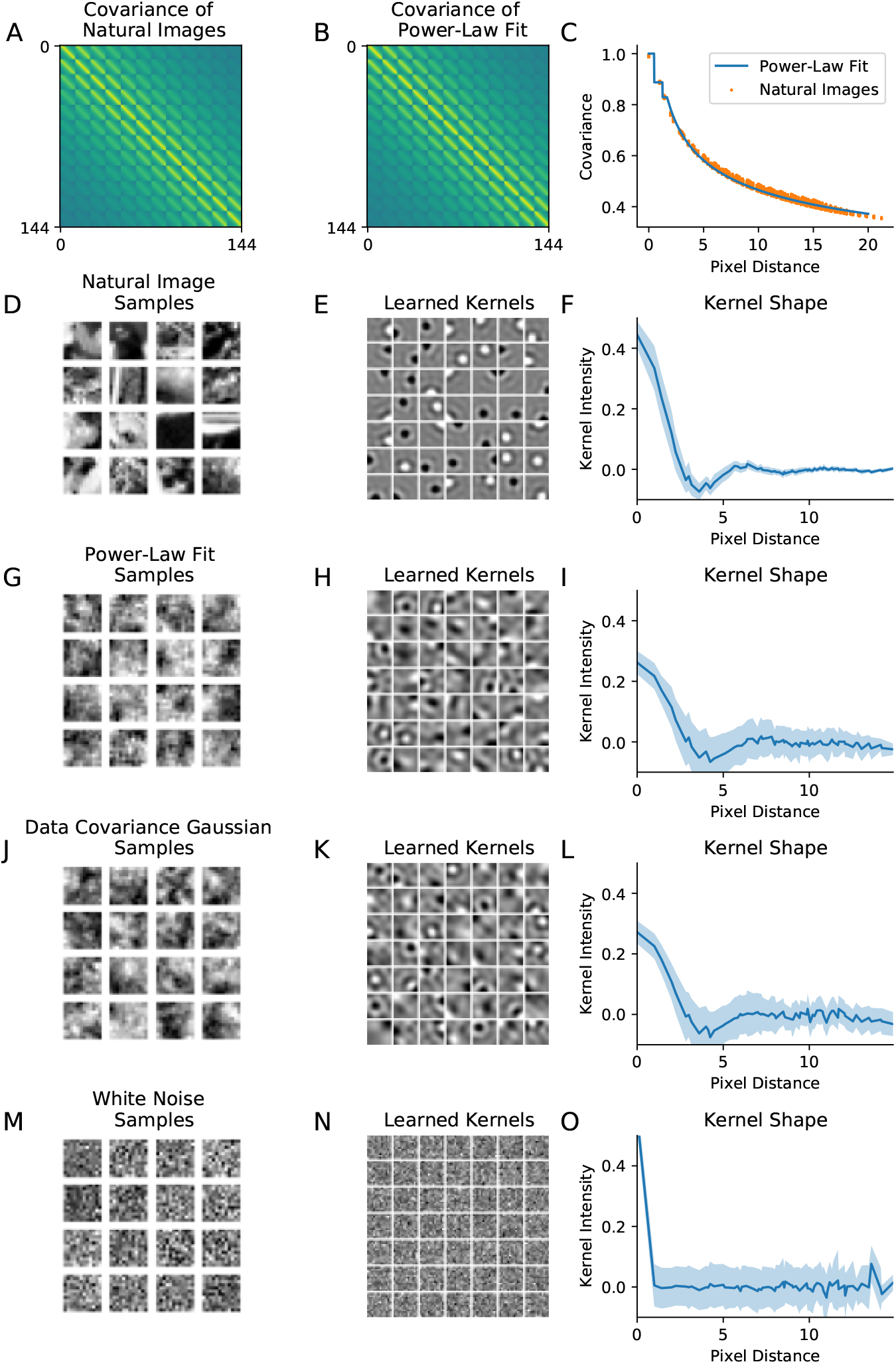
Learned kernels do not converge to circular center-surround shapes for non-natural image distributions. **A.** Covariance matrix of the natural images, calculated from 100,000 random image patches from Doi and Lewicki (2007). **B.** Covariance matrix constructed from the power-law fit of the values in **A**. **C.** Covariance of the 100,000 natural image patches (orange dots) and the power-law fit of the covariance (blue curve) defined as: *C_ijki_* = min(1,0.9765*r*^−0.322^) where 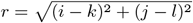. **D.** Natural image samples. **E.** Learned kernels when natural image samples such as **D** are the inputs to the model. **F.** Mean (blue curve) and standard deviation (light blue shade) of the kernel intensities as a function of the pixel distance from the location of the peak intensity (regardless of the ON and OFF centers). **G.** Image samples drawn from a multivariate Gaussian distribution with zero mean and the power-law fit covariance in **B**. **H.** Learned kernels when samples as in **G** are used as inputs. **I.** Same as **F** but with the kernels in **H**. **J.** Image samples drawn from a multivariate Gaussian with zero mean and the empirical covariance of natural images in **A**. **K.** Learned kernels when samples from **J** are used as inputs. **L.** Same as **F** but with the kernels in **K**. **M.** White noise image samples where each pixel is drawn i.i.d. from the standard normal distribution. **N** Learned kernels when the white noise samples such as those in **M** are the inputs of the model. **O.** Same as **F** but with the kernels in **N**. Only in the case of natural images do center-surround kernels emerge from training.

**Supplementary Figure 8.**
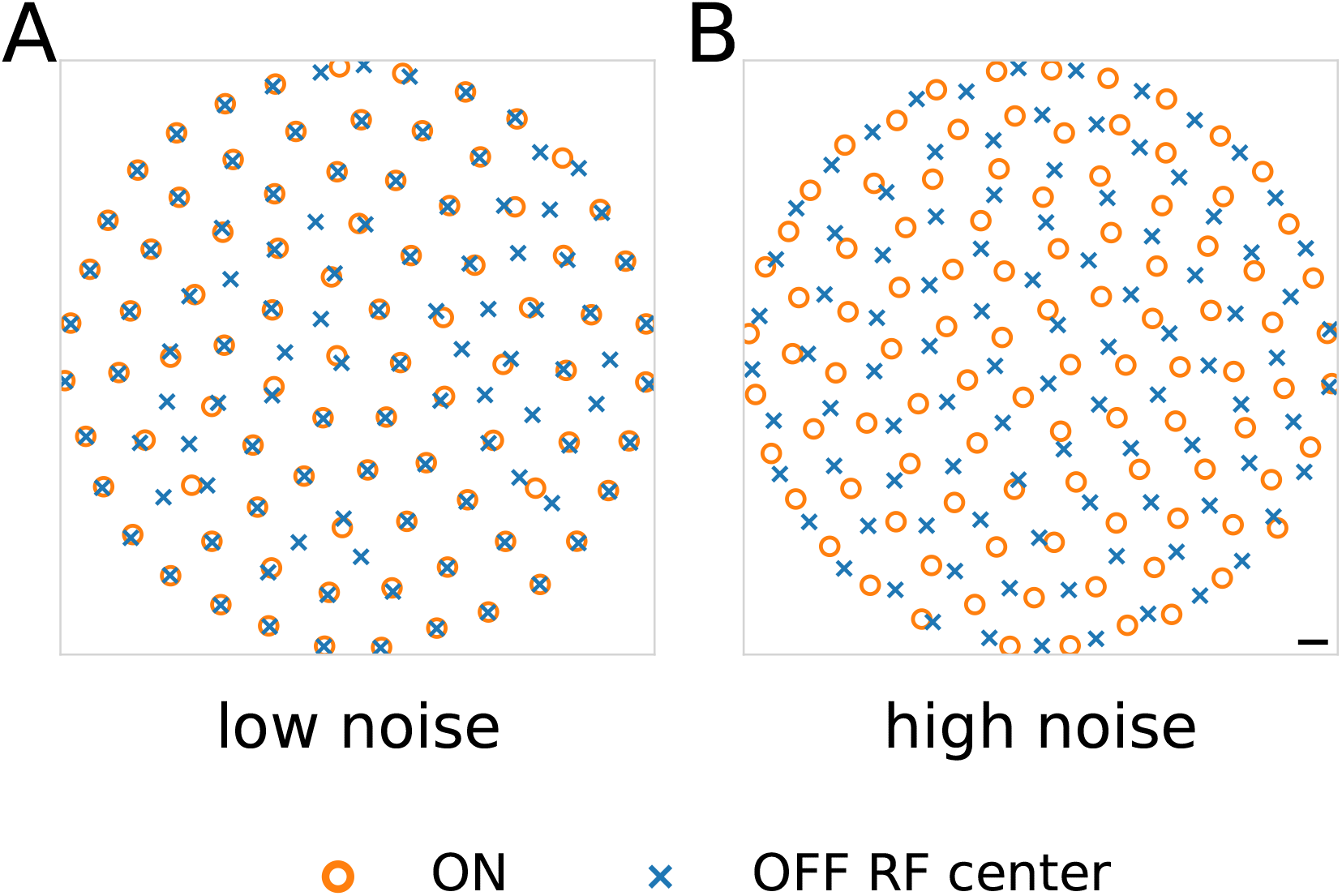
Larger image patches also produce anti-aligned mosaics. Receptive field centers for ON (orange circles) and OFF (blue Xs) cells under different noise configurations for patches of 25 x 25 pixels and 196 units. **A**: (*σ*_in_=0.4, *σ*_out_=1.0), **B**: (*σ*_in_=0.4, *σ*_out_=3.0). Noise parameters in **A** produce aligned mosaics, while those in **B** produce anti-aligned mosaics. Scale bar is width of one image pixel.

**Supplementary Figure 9.**
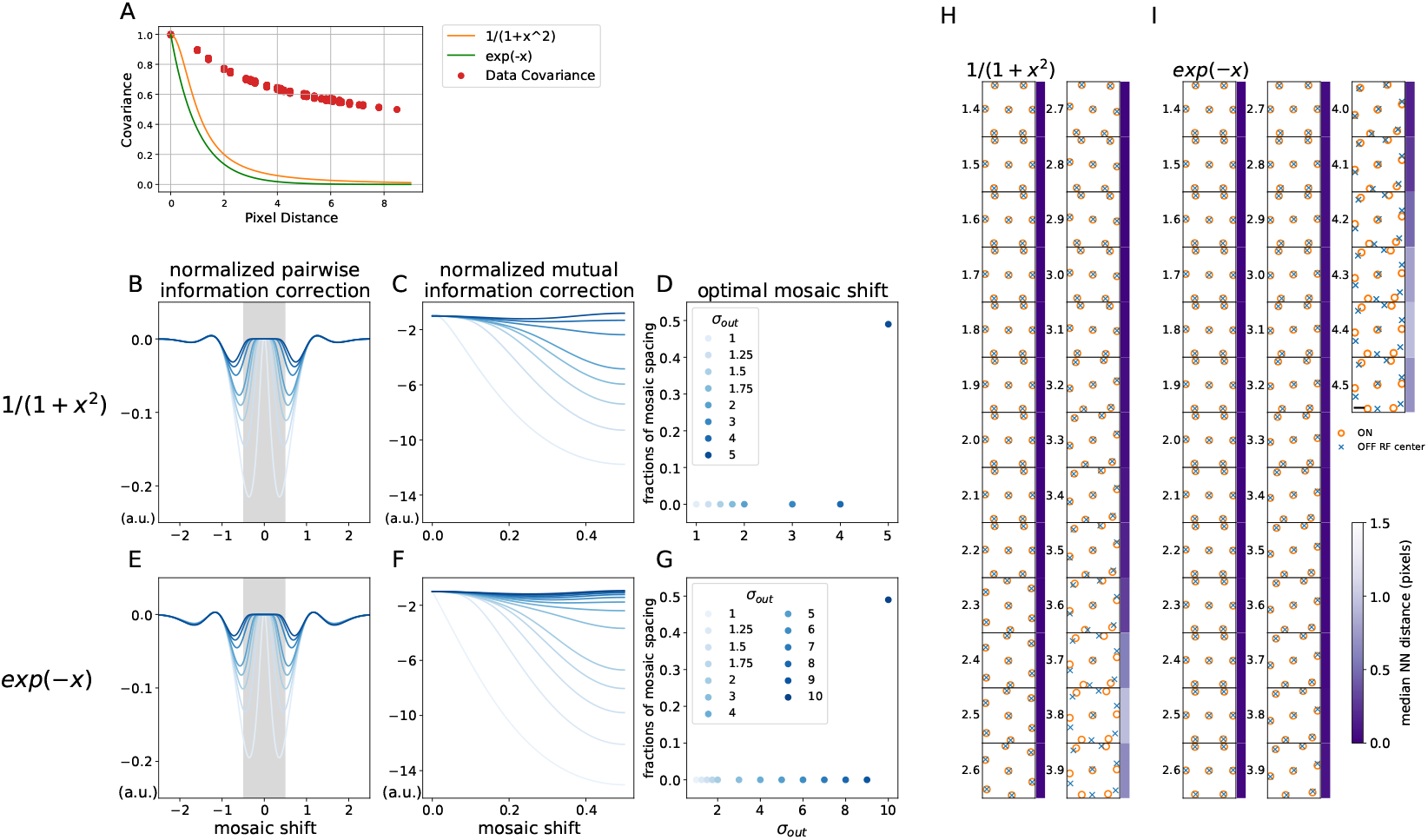
The transition to anti-aligned mosaics occurs with short-range correlations. **A.** Covariance of the 100,000 natural image patches (red dots) with long-range correlations, and two alternative correlation structures: 1/(1 + *x*^2^) (orange curve) and exp(-*x*) (green curve). **B-D.** Replication of **Fig 5A-C** when a model with assumed center-surround kernels is trained on images drawn from a multivariate Gaussian distribution with zero mean and the covariance function 1/(1 + *x*^2^). Unlike with natural images, anti-alignment occurs at very high levels of output noise. **E-G.** Same as **B-D** but when trained with the image samples drawn from a multivariate Gaussian distribution with zero mean and the exp(-x) covariance. **H.** Optimized ON and OFF receptive field mosaics of the 2D one-shape model using image patches with the 1/(1 + *x*^2^) covariance. The phase transition is observed around *σ*_out_=3.7, which is a higher noise than that producing the transition for natural images in **Fig 3. I.** Same as **H** but for exp(-x) covariance. The phase transition is observed around *σ*_out_=4.3, which is even higher than the case in **H**.

## II. MODELING DETAILS

### A. Mutual information in the linear-nonlinear model

We follow the conventions of Karklin and Simoncelli (2011). Let **x** ∈ ℝ^*D*^ be a vectorized representation of an input image. We assume *J* neurons characterized by linear filters **w**_*j*_, (||**w**_*j*_|| = 1) for *j* = 1…*N*, which form the columns of a matrix **W** ∈ ℝ^*D*×*J*^. We assume pixelwise input noise to each image 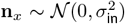 and output noise in firing rates 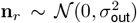 so that the random input image and firing rate are **X** = **x** + **n**_*x*_ and **R** = **r** + **n**_*r*_, respectively. With these conventions, the firing rate for the *j*th neuron given input **x** is

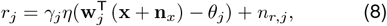

with *γ_j_*, the neuron gain, *θ_j_* its response threshold, and *η* the softplus nonlinearity described in Methods. Note that, in principle, this firing can be negative for sufficiently large and negative *n_r,j_*. As in Karklin and Simoncelli (2011), we notheless treat **Eq 8** as a useful approximation.

Under these assumptions, the covariance matrix of firing rates given input image is

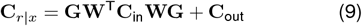

with 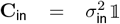, 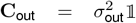, and **G**(**x**) = diag(*γ,η*(**w**^⊤^(**x** + **n**_*x*_) – *θ*)) a matrix of derivatives of the nonlinearities. As in Karklin and Simoncelli (2011), we make a local Gaussian approximation to the image distribution with covariance **C**_*x*_, allowing us write the posterior *p*(**x**|**r**) as Gaussian with covariance

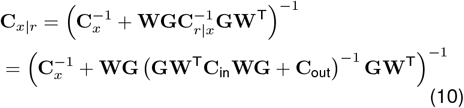

and the conditional (differential) entropy as

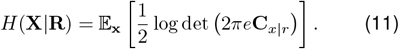

Since *H*(**X**) is an unknown constant, maximizing the mutual information 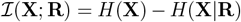 involves *minimizing* **Eq 11** subject to a cost per spike that sets constrains the mean firing rate of each neuron:

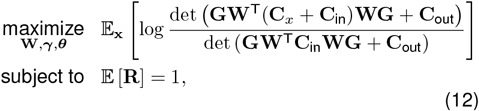

where the optimization objective follows from **Eqs 10** and **11** via the Matrix Inversion Lemma and we implement the constraint using an augmented Lagrangian method as detailed in Methods. Finally, the following rewriting of **Eq 12** will be useful, particularly in our analysis of the single-neuron case:

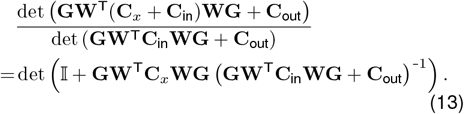

### B. Analytic model of alignment in 1-D

To better understand the optimization problem posed in **Eq 12**, particularly the contributions of kernel shape, image statistics, noise, and response thresholds to the transition from aligned to anti-aligned mosaics, we consider a simplified model that nonetheless exhibits the same behavior. In particular:

1. We restrict ourselves to a one-dimensional space in which ON and OFF mosaics are assumed to be an equidistant array of receptive field centers (distance = 1) for both mosaics. The ON cells are located at positions 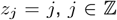, and the mosaics are allowed to rigidly shift relative to each other, so that OFF cells are located at 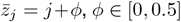.
2. We assume a fixed kernel shape for all neurons, meaning that receptive fields differ only in their center locations: *w_j_*(*z*) = ±*κ*(|*z* – *z_j_*|) for a kernel shape function *κ*(*z*), where *z_j_*, is the location of *j*-th neuron’s kernel center.
3. We likewise assume identical nonlinearities for all units: *f_j_*(*y*) = *γ*(*y* – *θ*)_+_, where *γ* is the gain, *θ* the response threshold parameter, and (·)_+_ = max(·, 0) denotes positive rectification.
4. Images **x** are assumed to follow a multivariate Gaussian distribution 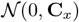, where the covariance between locations *z_i_* and *z_j_*, only depends on |*z_i_* – *z_j_*|, defined as a function *C_x_*(*z*). In analogy with the finding that the amplitudes of natural images decrease as ~ 1/*k* with *k* the spatial frequency (power spectral density ~ 1/*k*^2^), for our numerical examples, we likewise assume a *one-dimensional* scale-free distribution 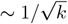 (power spectrum ~ 1/*k*). As shown in **Fig S5B**, the phase transition still takes place even with this Gaussian image assumption.

Now, for a pair of neurons *j* and *k*, the scalars 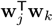 and 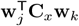 depend only on |*z_j_* – *z_k_*|, the distance between their centers, which we write in the frequency domain as:

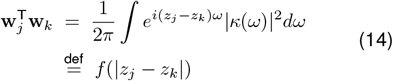

**Supplementary Figure 10.**
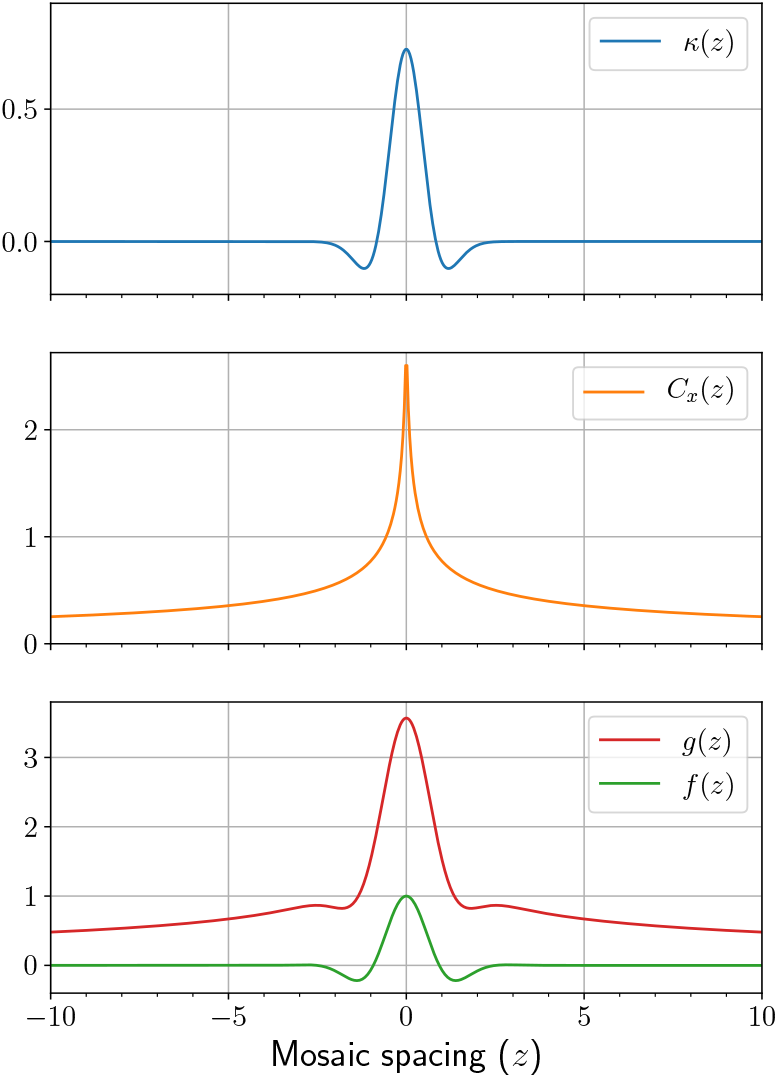
Plots of *κ*(*z*), *C_x_*(*z*), *f*(*z*), and *g*(*z*) using trained parameters from a one-shape model and an approximation of the data covariance: *κ*(*z*) ∝ *e*^−1.76*z*2^ – 0.65*e*^−1.12*z*2^ and 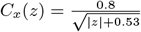.

and

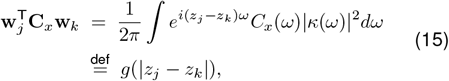

where *C_x_*(*ω*) and *κ*(*ω*) are Fourier transforms of *C_x_*(*z*) and *κ*(*z*), respectively. Using any **w**, we can define the constants *f*_0_ and *g*_0_ for the special case where *z_j_*, = *z_k_*:

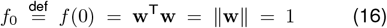

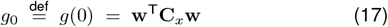

where we used our assumption that the kernel **w** is a unit vector in **Eq 16**. An example of these functions is shown in **Sup Fig 10**.

#### The single-neuron case: 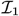

We first consider the mutual information capacity of a single neuron in isolation. Since we have assumed all units to be identical, the same expression holds for all def ON and OFF cells, and we let 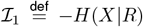 in this single-channel case. From **Eq 13**, this quantity is

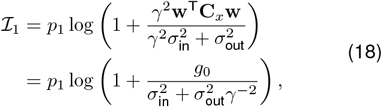

where 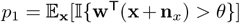 is the probability that the neuron is active, i.e., that **w**^⊤^(**x** + **n**_*x*_) > *θ*. Now, since **x** + **n**_*x*_ is Gaussian with mean 0 and covariance **C**_*x*_ + **C**_in_, we have 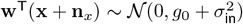 and

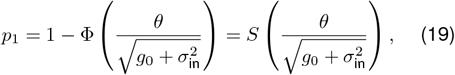

where Φ and *S* are the cumulative distribution function and the survival function of the standard normal distribution, respectively. Note that if we define 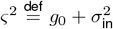 to denote the signal variance and 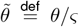 to denote the effective threshold, we see that the effective threshold increases when signal variance decreases. We can calculate *γ* as:

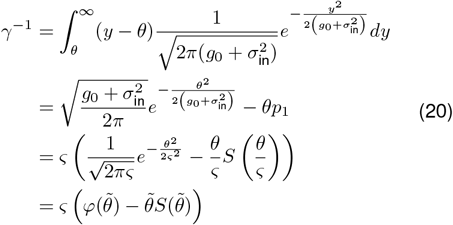

where *φ* is the probability density function of the standard normal distribution. Substituting this to **Eq 18,** we get:

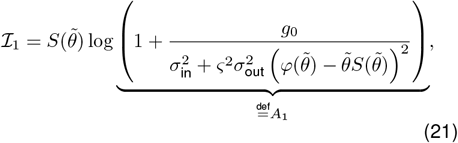

which was used to plot the single-neuron mutual information contributions in **Fig 4A.**

#### A single-neuron case with heavy tails

To investigate the relationship between threshold and outliers in the image distribution, we performed the analysis in **Fig 4B,** which replaced the Gaussian image distribution with a Student’s *t*-distribution. This allowed us to adjust the heaviness of the tails of the distribution by changing the degrees of freedom from *ν* = 4 (heavy tails) to *ν* = ∞ (Gaussian). In this case, a formula similar to **Eq 21** can be derived in a similar manner to **Eq 20**.

More explicitly, if **x** ~ *t_ν_* (0, **C**_*x*_), then **w**^⊤^(**x** + **n**_*x*_)/*ς* is likewise *t_ν_*-distributed, and we have:

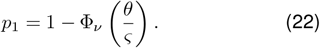

where Φ_*ν*_ is the cumulative distribution function of the *t*-distribution. Moreover,

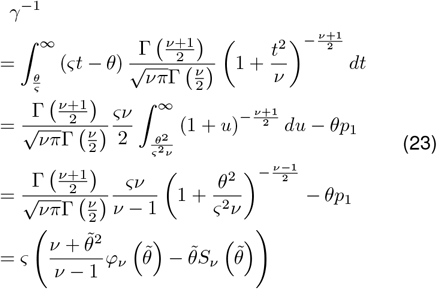

where *φ_ν_* and *S_ν_* are respectively the probability density function and the survival function of Student’s *t*-distribution with *ν* degrees of freedom. We see that 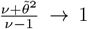 as *ν* → ∞, in which case **Eq 23** is identical to **Eq 20**. These results can then be substituted into **Eq 18** to derive an expression analogous to **Eq 21**.

#### Decomposition of the mutual information

In previous sections, we have considered the encoding properties of a single neuron in isolation. Here, we return to the full optimization objective in **Eq 12,** showing how we can rewrite this as a sum of independent, single-neuron contributions plus pairwise (and higher-order) correction terms.

We begin by noting that the expectation in **Eq 12** over images **x** involves a sum over patches that only activate subsets of neurons. More specifically, with the hard rectifying nonlinearity assumed in the previous section, we have 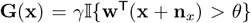 for ON cells and the same for OFF cells with *θ* → −*θ*. Here, 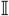 is the indicator function, which is 1 when its argument is true and 0 otherwise. Thus, the gain matrix is diagonal, with entries equal to either 0 or *γ*, and the matrix products involving **G**(**x**) in **Eq 12** are thus low-rank for most **x**. As a result, we can rewrite the expectation as a sum over log determinants of low-rank matrices, each of which corresponds to a specific collection of active neurons.

**Supplementary Figure 11.**
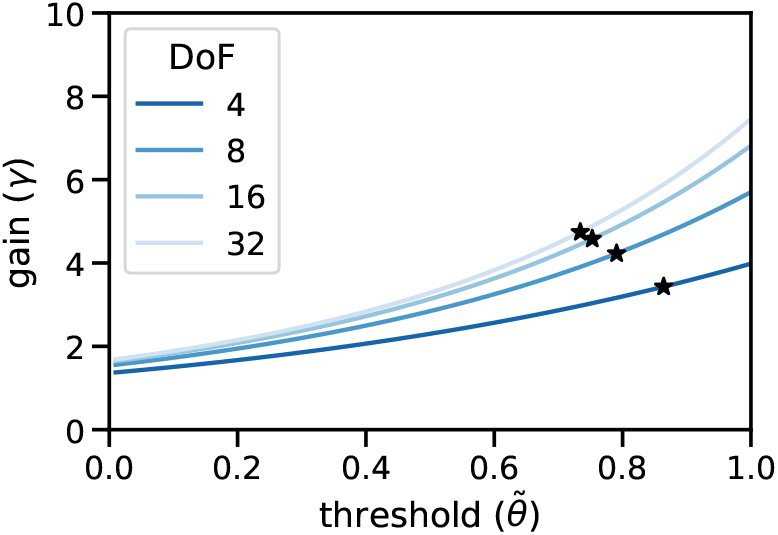
When the input distribution has heavier tails, modeled with Student’s t distributions with varying degrees of freedom, the optimal gain decreases with heavier tails. The optimal thresholds are depicted in **Fig 4B,** and optimal gains are computed using the optimal thresholds and **Eq 23** with *ς* = 1.249.

For example, consider a three-neuron system, whose activity can be coded as a binary triple, (e.g., 010). Let *A* be the determinant in **Eq 13**. Letting *p*_(triplet)_ denote the probability that each subset is activer over the image set and *A*_(triplet)_ be the corresponding determinant ratio, the expectation can be expanded as:

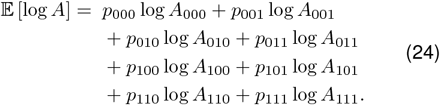

Unfortunately, even for a Gaussian image distribution, calculating *p* for a specific triple involves a highdimensional Gaussian integral bounded by a triple of hyperplanes in image space. However, we can rewrite this collection of terms as:

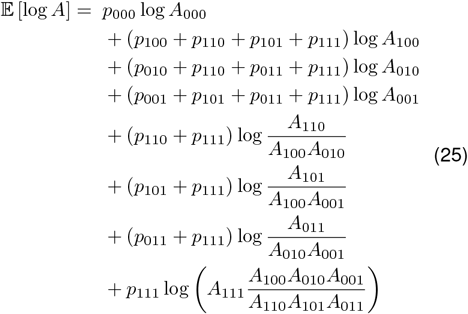

The first term corresponds to the case when no neuron is active, for which log *A*_000_ = 0. For the next three terms, each of the probabilities in parentheses is a *marginal* probability of a neuron being active. That is, it is equal to *p*_1_ as calculated above. Similarly, the next three terms constitute pairwise corrections to the first set of terms: they involve marginal probabilities of pairwise activation, and their log ratios vanish when the individual neurons are independent (e.g., *A*_110_ = *A*_100_*A*_010_). Thus, as the distance between a pair of receptive field centers increases beyond the image correlation length scale, these correction terms should vanish.

In our analysis, we focus on the limit in which only first- and second-order interactions are non-negligible. This is an assumption that does not, in fact, hold for natural images (or even our Gaussian images with long-range correlations), though as we shall see, these interactions are sufficient to demonstrate the existence of a phase transition. Moreover, as Figures 5 and 6 show, the same phase transition occurs in simulation when images with long-range correlations are used, suggesting that the principles we identify in the pairwise limit generalize to the full model.

#### *Pairwise interactions: h*_2_ and *h*_2′_

Returning to our 1-D model with *N* ON and *N* OFF neurons, we note that, as per **Eq 5**, we can further decompose the pairwise corrections into two terms, one corresponding to ON-ON and OFF-OFF corrections, the other to ON-OFF pairs. More explicitly, if we denote by *a*, and *ā_j_*, the binary variables corresponding to whether the ON cell at location *j* and the OFF cell at location *j* + *ϕ* are active, respectively, we can write the mutual information up to pairwise corrections as:

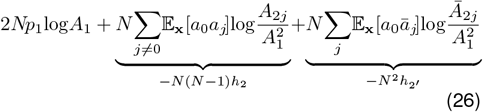

where the first term contains 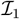 from **Eq 21**, and the next two terms correspond to pairwise interactions between neurons of the same and different polarities, respectively:

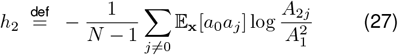

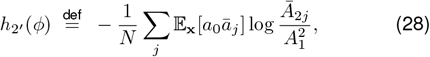

Note here that we have used translational invariance to write the pairwise corrections as the interaction between a single ON cell at 0 and all other cells and that there are 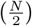 such pairs. Likewise, since the ON and OFF cells are identical up to a polarity flip, the *h*_2_ term includes both mosaics’ self-interactions for a total of *N*(*N* – 1) such terms. Similarly, there are *N*^2^ ON-OFF interactions, resulting in the differing normalizations above.

Clearly, only the last term in **Eq 26** is a function of *ϕ*, and we only need to consider the value *ϕ* that maximizes *h*_2_/ to determine whether alignment (*ϕ* = 0) or anti-alignment (*ϕ* = 0.5) is preferred. We know from **Eq 18** that

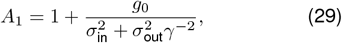

and we can calculate the two-neuron determinant ratio 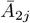, as follows: First define matrices **P** and **Q** from **Eq 12** to write

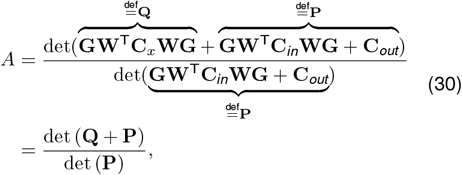

then substitute kernels **w**_0_ and – **w**_*j*_, (negative because of the opposite polarity):

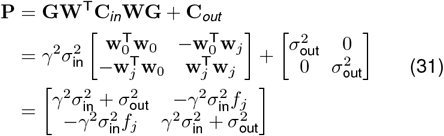

and

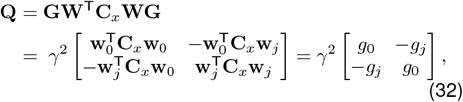

where we define 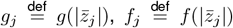, and thus *f*_0_ = 1. We then have

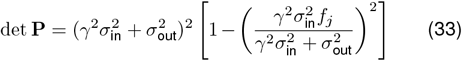

and

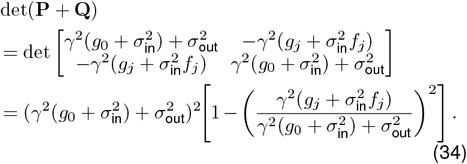

**Supplementary Figure 12.**
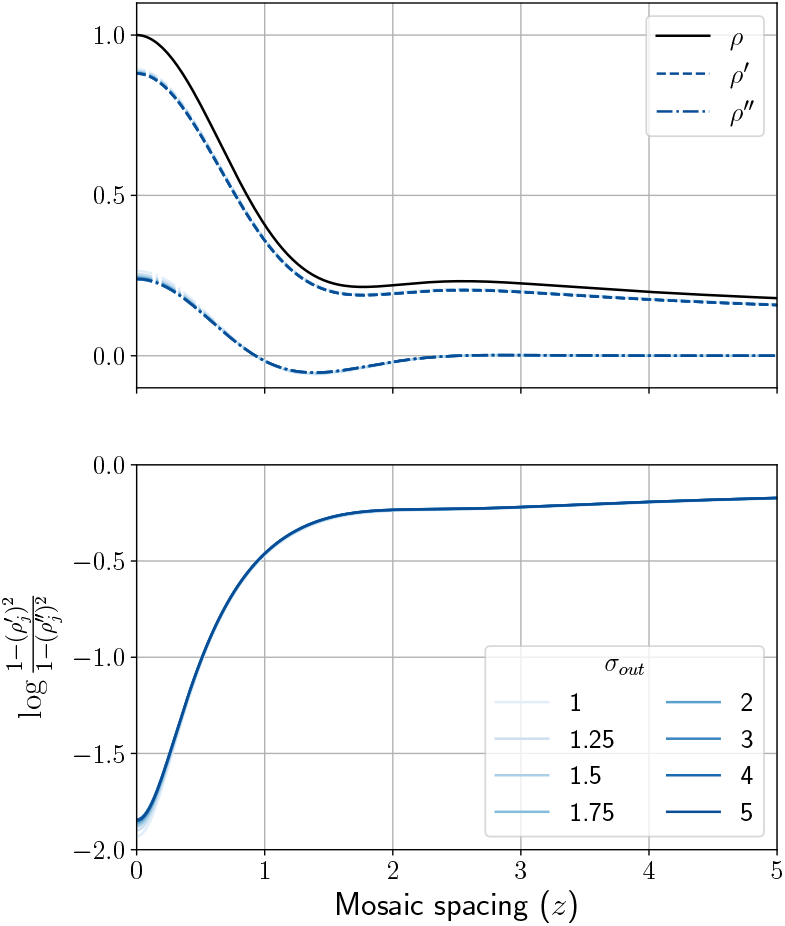
Plots of the correlation-like quantities *ρ, ρ′*, and *ρ”* as defined in **Eqs 39** and **35** and the log ratio 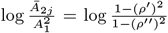, with respect to color-coded *σ*_out_ values. These depend very weakly on *σ*_out_, suggesting the 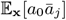 term in **Eq 28** plays a more crucial role in the phase transition.

Collecting these with the expression for *A*_1_ in **Eq 29**,

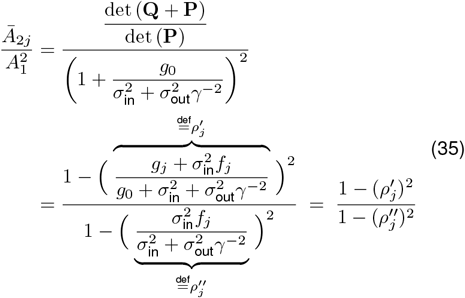

Note that since *f_j_* ≤ 1 by definition and *g_j_* ≤ *g*_0_ for most reasonable image distributions, both terms in brackets are less than 1. Also, as *j* → ∞, we have *g_j_, f_j_*, → 0, so the term vanishes asymptotically. In fact, *f_j_*, vanishes very quickly (as soon as the receptive fields have negligible overlap), while *g_j_*, diminishes much more slowly, since **C**_*x*_ contains long-range correlations. Note that *f_j_*, and *g_j_*, are the only terms here that depend on *ϕ*. More importantly, this term only varies weakly with *σ*_out_ as shown in **Sup Fig 12**, suggesting it does not play a crucial role in the phase transition.

Now, as for the coactivation probability 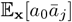, we consider 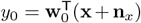 and 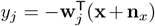 whose joint distribution follows

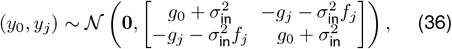

which means that 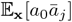 can be calculated viaaGaussian integral over the region *y*_0_ > *θ* and *y_j_*, > *θ*:

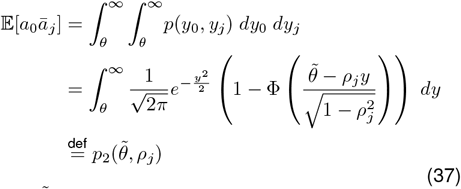

where 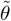 and *ρ_j_*, are defined as

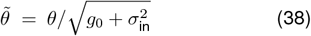

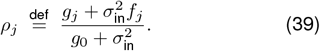

Putting all this together, we can write *h*_2′_ as

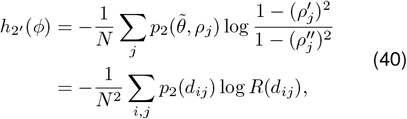

where again, the dependence on *ϕ* is solely through *f_j_*, = *f*(|*j* + *ϕ*|) and *g_j_*, = *g*(|*j* + *ϕ*|) and we have used translation invariance to rewrite these arguments as *d_ij_*, the distance between ON-OFF pairs. Finally, since *f_j_* ≤ 1 by definition, 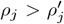, always, and 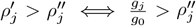 (which holds empirically with our image distribution and optimized kernels), we have 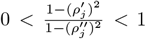 and thus 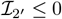.

For the results plotted in **Fig 5A**, we numerically integrated **Eq 37** to calculate *p*_2_ and used kernels and covariance parameters taken from a trained one-shape model, shown in **Sup Fig 10**, to calculate **C**_*x*_, *f_j_* and *g_j_*. To make these quantities comparable as *σ*_out_ increases (and thus *p*_2_ decreases), **Fig 5A** plots the normalized contribution 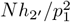.

### C. Redundancy in aligned and independent ON-OFF pairs

We have argued in **Fig 4D-E** that redundancy is reduced by aligning a single OFF cell (in the case of low *θ*) and anti-aligning multiple OFF neighbors (for high *θ*) inside a “reduced redundancy” region of small *p*_2_ around each ON cell. However, this characterization may be objected to on the grounds that, in fact, aligned response fields are highly redundant, since the activation of one by a stimulus excludes activation of the other.

In a limited sense, this is true. Consider replacing our monotonically increasing nonlinearity *h* with a simple threshold nonlinearity, such that neurons fire at a constant rate when active, similar to Gjorgjieva *et al.* (2014). In this case, we can calculate for a single neuron with *θ* = 0

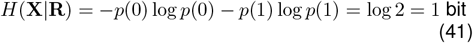

since the neuron is expected to be active half the time. Clearly, for two independent cells (e.g., those with receptive field centers separated by a distance much larger than the image correlation scale), we then expect 2 bits of conditional entropy. However, for an aligned ON-OFF pair, we would have

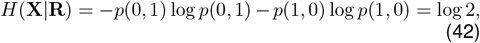

just as before. Thus, when each neuron has a capacity of one bit, two aligned receptive fields are completely redundant.

However, the assumption of a graded firing rate conveys additional information about the stimulus beyond its sign, and this information is not fully shared even by aligned receptive fields. For instance, in the case of a single neuron, we have from **Eqs 21** and **29**

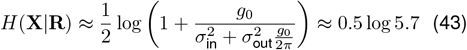

for *θ* = 0, *σ*_in_ = 0.4, *σ*_out_ = 1, *g*_0_ ≈ 3.8. However, this is *differential* entropy. To relate this to the answer in the 1-bit case, we need to consider a quantization of the signal. If we assume that the signal can be quantized using 1 bit for its sign and *n* bits for its magnitude, we have, for *n* sufficiently large Cover (1999)

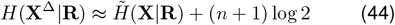

where **X**^Δ^ is the quantized variable, and we denote the differential entropy by 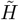.

From this result, it is clear that, while far-separated receptive fields encode precisely 2*H*(**X**^Δ^|**R**), perfectly aligned receptive fields encode only 2*H*(**X**^Δ^|**R**) – log 2, where the correction is due to the shared sign bit. Thus, while it is correct that aligned receptive fields do exhibit some redundancy (1 bit for *θ* = 0, less as *θ* increases), this is typically much smaller than the information about stimulus magnitude. More importantly, when considering how to align ON and OFF cells, informational losses due to shared sign bits are dwarfed by the size of *N*^2^*h*_2_ for neighbors located outside the gray zone in **Fig 4A.**

